# Clusterin regulates microglial inflammation and cognitive function independent of amyloid pathology in Alzheimer’s disease

**DOI:** 10.64898/2026.07.30.741791

**Authors:** Punam Rawal, Hee-Jung Moon, Vanessa Nguyen, Sophia Khatri, Melissa A. Larson, Jay L. Vivian, Takaomi C. Saido, Liqin Zhao

## Abstract

Clusterin (CLU) is a major genetic risk factor for late-onset Alzheimer’s disease (AD), yet the mechanism underlying this risk remains unclear. This study investigated the role of CLU in regulating microglial inflammation, amyloid pathology, and cognitive function in CLU-deficient and AD models. CLU synthesis and secretion in microglia were highly dynamic, with minimal expression at rest but markedly increased expression upon pro-inflammatory stimulation. CLU deficiency amplified microglial activation and inflammatory responses, whereas introducing recombinant CLU protein reduced microglial inflammation. In a human mutant APP knock-in AD mouse model, loss of CLU impaired learning and memory despite a reduced amyloid burden. snRNA-seq analysis further revealed that CLU loss disrupted the balance between excitatory and inhibitory neurons. These findings provide novel insights indicating that CLU likely functions as a negative feedback regulator of microglial activation and inflammation, thereby promoting microglial resolution and neuronal homeostasis, and consequently, cognitive outcomes independent of amyloid pathology.

## INTRODUCTION

Clusterin (CLU), also known as apolipoprotein J (ApoJ), is currently recognized as the third most significant genetic risk factor for late-onset Alzheimer’s disease (AD) (*1–4*). The compelling link between CLU and AD is strongly supported by clinical findings from examinations of postmortem brain tissue, cerebrospinal fluid (CSF), and plasma. For instance, elevated levels of CLU were found in the hippocampus and frontal cortex of AD-afflicted brains; however, no significant changes were detected in the cerebellum of AD brains and brains of vascular dementia (VaD) (*5*). Moreover, there was no correlation between CLU expression and the extent of amyloid β (Aβ) plaque deposition, suggesting that the increased CLU in AD brains was likely independent of amyloid pathogenesis (*5*). Furthermore, in a study focused on the relationship between CLU expression and formation of neurofibrillary tangles (NFT), CLU was found to be predominantly expressed in NFT-free cortical neurons localized in brain regions most vulnerable to AD, whereas it was nearly undetectable in NFT-expressing neurons in the same regions, pointing to the hypothesis that increased expression of CLU in AD brains might represent a protective response in promoting neuronal survival in a toxic environment (*6*). In the CSF, CLU was shown to interact with soluble Aβ, suggesting that CLU may play a role in Aβ transport through the blood-brain barrier (*7, 8*). In the plasma, a study using discovery-phase proteomics to identify proteins correlating with AD pathology revealed a strong association between CLU levels and entorhinal cortex atrophy, baseline disease severity, and rapid clinical progression (*9*). Additionally, plasma CLU levels positively correlated with the rate of brain atrophy, while CLU expression in AD-vulnerable brain regions correlated with mild cognitive impairment (MCI) patients compared to control groups (*10, 11*). Despite the wealth of evidence demonstrating the link of CLU to AD, the precise pathophysiological role of CLU in AD remains elusive (*4, 12*). The immunomodulatory role of CLU has been explored in peripheral tissues (*13, 14*); however, whether this immunomodulatory role of CLU extends to the brain remains poorly understood.

Recent genome-wide association studies (GWAS) have identified a number of single-nucleotide polymorphisms (SNPs) in genes associated with an increased AD risk, many of which are predominantly or exclusively expressed in microglia (*15–18*). This highlights the essential role of microglia-mediated immune responses as a critical regulator of AD pathology, which is further corroborated by gene regulatory network analyses in postmortem human brains (*19*). Microglia, as brain-resident immune cells, function as phagocytes, defending the brain through the release of cytokines and the phagocytosis of cellular debris and aggregated proteins (*15*). While proinflammatory cytokines are beneficial and support healing in acute injury, they can become detrimental under chronic conditions, disrupting repair and promoting degeneration (*20*). Thus, it becomes apparent that mechanisms that properly regulate microglial inflammation and restore microglial homeostasis likely play a pivotal role in maintaining brain health and reducing the risk of AD (*21*).

This study investigated the immunomodulatory role of CLU in the brain, with a focus on microglia. Using CLU-deficient (*Clu*^−/−^) and WT (*Clu*^+/+^) mice, as well as primary and immortalized microglial cultures, CLU deficiency was associated with elevated microglial activation and inflammation. Although microglia exhibited low CLU levels under resting conditions, CLU expression and secretion were significantly upregulated following stimulation with lipopolysaccharide (LPS). Treatment with recombinant CLU (rCLU) protein attenuated LPS-induced microglial activation and restored cellular morphology, supporting a suppressive role for CLU in microglial inflammation. In the AD mouse model harboring *App*^NL-F/NL-F^, loss of CLU resulted in reduced Aβ plaque burden but impaired cognitive function. Single-nucleus RNA sequencing (snRNA-seq) revealed a disruption in the balance between excitatory and inhibitory neurons, marked by a significant loss of excitatory neurons. These findings support the conclusion that CLU plays an important role in brain health by regulating microglial inflammation and neuronal homeostasis, thereby influencing cognitive outcomes independent of Aβ pathology.

## RESULTS

### Microglia expressed low levels of CLU under resting conditions

Astrocytes have been widely recognized as a significant source of CLU in the brain (*4, 22, 23*). However, studies investigating the expression of CLU in microglia are limited and have yielded conflicting findings (*22, 24*). Therefore, our initial aim was to perform an in-depth analysis of microglial CLU synthesis and secretion profiles. The identification and purity of primary astrocytes, primary microglia, and immortalized microglial (IMG) cells were validated by immunocytochemistry and by expression of specific protein markers (Fig. 1A and Fig. S2). The *Clu* mRNA levels in primary astrocytes, primary microglia, and IMG cells were determined using RT-qPCR (Fig. S1). As expected, the cycle thread or Ct value obtained from astrocytes was found to be substantially lower compared to both primary microglia and IMG cells, confirming a much higher expression of *Clu* mRNA in astrocytes compared to microglia.

**Fig. 1.**
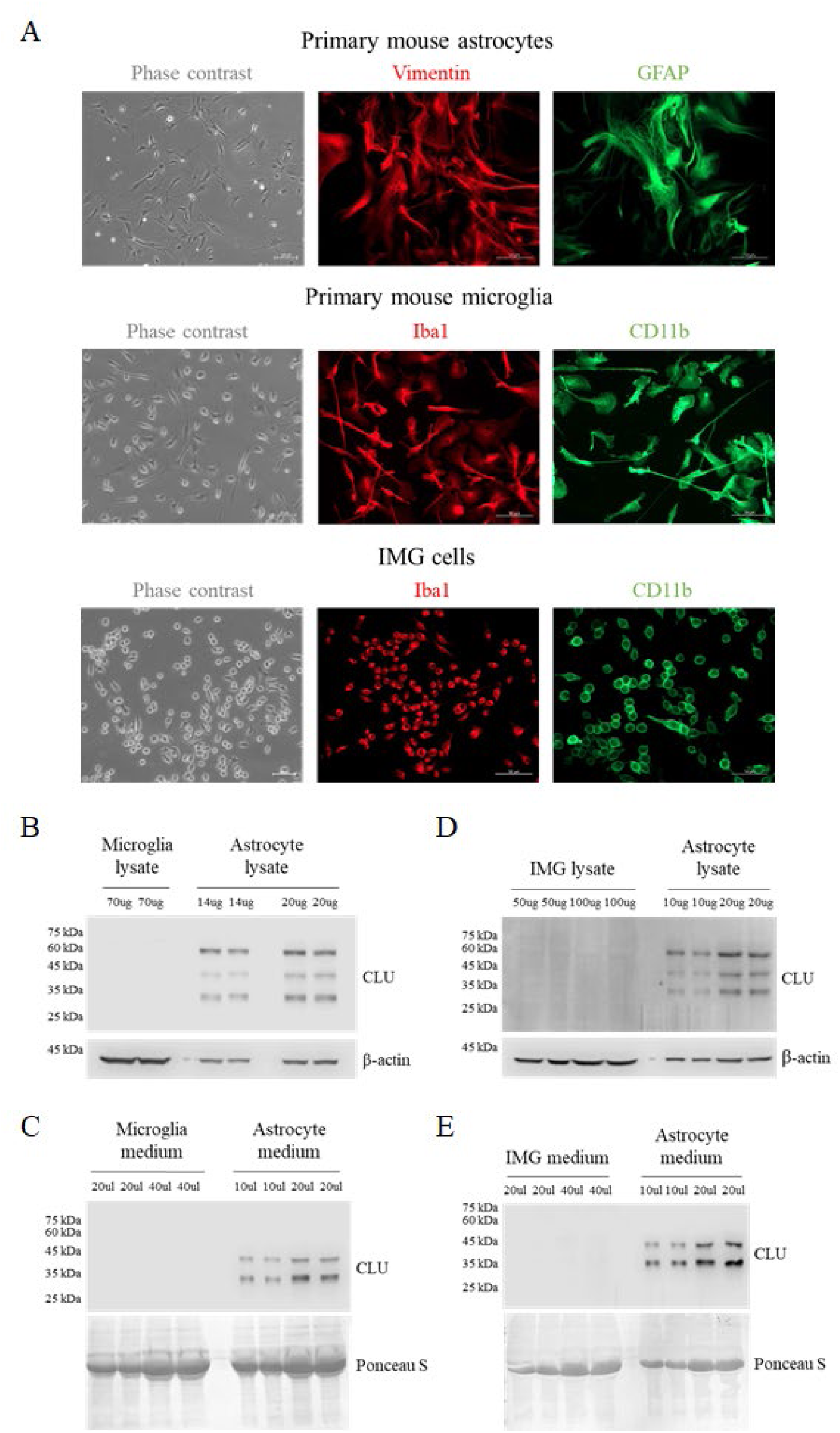
Characterization of astrocytic and microglial CLU expression under resting conditions. **(A)** Validation of cell models used in the study. Primary mouse astrocytes are confirmed by cell morphology and prominent Vimentin and GFAP expression. Primary mouse microglia and IMG cells are confirmed by cell morphology and prominent Iba1 and CD11b expression. **(B-E)** CLU protein expression and secretion in microglia were compared with those in primary astrocytes: (B) primary microglia lysate and (C) medium, (D) IMG cell lysate and (E) medium. Given the relatively low microglial *Clu* mRNA level (Fig. S1), the protein load for microglia samples was higher than that for astrocytes.

Building upon this mRNA data, the protein expression profile of CLU in both cell lysates and medium fractions was determined. Given the low levels of *Clu* mRNA in microglia, a higher amount of protein was loaded for microglia samples relative to astrocytes. Notably, the Western blot analysis successfully detected all isoforms of mature CLU in astrocytes (Fig. 1B-E). This included the high mannose modified intermediate form at 60 kDa in the cell lysate, as well as the two subunits of mature CLU (alpha subunit at 34-37 kDa and beta subunit at 36-39 kDa) in both the lysate and medium fractions (*25*). However, in the case of microglia, a minimal, barely detectable CLU expression was observed in the primary microglia cell lysate (Fig. 1B), primary microglia medium (Fig. 1C), IMG cell lysate (Fig. 1D), and IMG medium (Fig. 1E). These results indicate that under physiological resting conditions, microglia exhibit very low levels of CLU synthesis and secretion.

### Expression and secretion of CLU were markedly increased in microglia under inflammatory conditions

After observing low levels of CLU expression in microglia under normal conditions, the next aim was to examine the impact of inflammation on CLU expression. To achieve this, IMG cells were treated with LPS, a potent inducer of microglial inflammation (*26*). The cells were incubated with 10 ng/mL LPS or vehicle for 1 hour. Following the incubation, *Clu* mRNA levels were quantified, revealing a significantly elevated level of *Clu* mRNA in microglia upon LPS stimulation (Fig. 2A). Consistent with the mRNA results, a significant increase in microglial CLU protein expression was also observed in both IMG cell lysate and culture medium upon treatment with LPS (Fig. 2B). Furthermore, various LPS doses and incubation times were tested, and the data revealed a dose-and time-dependent upregulation of microglial *Clu* mRNA (Fig. 2C). Similarly, CLU protein levels were found to be upregulated with LPS treatment in both IMG cell lysate (Fig. 2D) and IMG medium (Fig. 2E). In summary, microglia carried out a rigorous synthesis and secretion of CLU in response to LPS-induced inflammation.

**Fig. 2.**
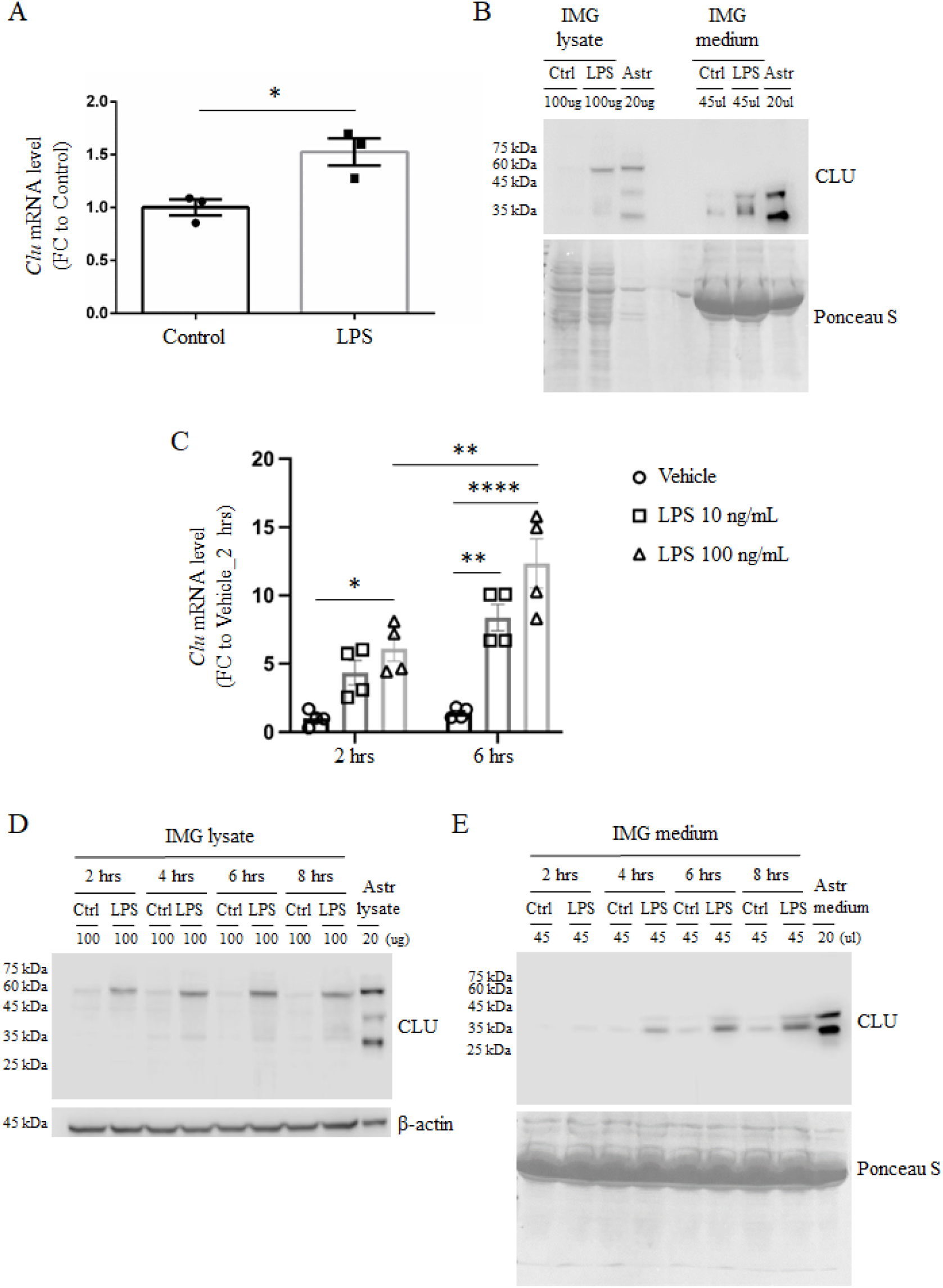
Microglial CLU expression and secretion were upregulated during LPS-induced inflammation. **(A)** IMG cells were treated with 10 ng/mL LPS or vehicle alone for 1 hour, after which cells were harvested for analysis of *Clu* mRNA levels by RT-qPCR. Results were normalized to the control group and compared using Student’s *t*-test. Data are presented as the group mean ± SEM. *n* = 3/group. **(B)** Consistent with mRNA levels, LPS treatment markedly increased CLU protein expression in both IMG cell lysates and medium fractions. **(C)** LPS induced time- and dose-dependent increases in Clu mRNA levels in IMG cells. Results were normalized to the vehicle + 2 hours group and compared using two-way ANOVA with Tukey’s post hoc test. Data are presented as the group mean ± SEM; *n* = 4/group. **(D-E)** LPS induced time-dependent increases in CLU protein expression in both IMG cell lysates and medium fractions. \**p* < 0.05, \*\**p* < 0.01, \*\*\*\**p* < 0.0001.

### CLU deficiency heightened microglial activation and inflammation in response to LPS stimulation

Microglia are highly dynamic cells, and their morphological phenotypes closely reflect their functional activity (*27*). Besides changes in the expression levels of various genes, microglial morphology can provide insights into their activation status (*28*). To address the role of CLU in microglial function and neuroinflammation, a novel mouse strain *(Clu*^−/−^) was generated for this study using CRISPR mutagenesis which lacked all CLU coding sequences and disrupted all CLU isoforms (Fig. 3A and Fig. S5). Primary microglial cultures derived from WT and novel *Clu*^−/−^ mice were stained with Iba-1 and compared for morphological differences (Fig. 3B). WT microglia were found to be extensively ramified with relatively long processes, indicative of an immunosurveillance state of microglia responsible for maintaining homeostasis of the neural environment (*29*). In comparison, *Clu*^−/−^ microglia displayed significantly shortened processes consistent with an activated state (Fig. 3C).

**Fig. 3.**
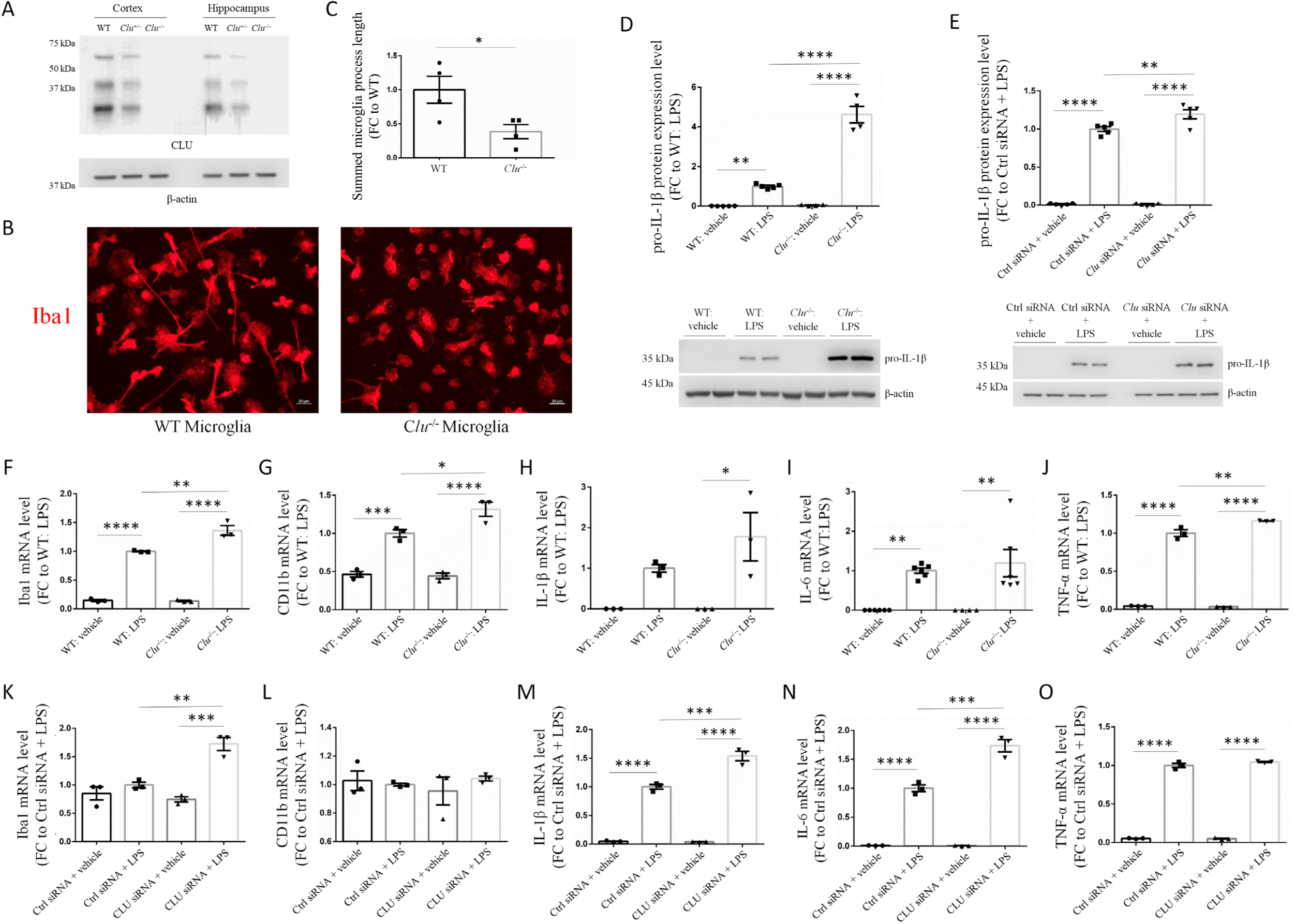
CLU deficiency induced an amoeboid-like microglial morphology and heightened microglial activation and inflammation in response to LPS stimulation. **(A)** Validation of the novel *Clu*^−/−^ mouse model generated in the present study; as expected, CLU expression was significantly reduced in *Clu*^+/−^ mice compared to WT mice, and no CLU expression was detected in *Clu*^−/−^ mice. **(B)** WT and *Clu*^−/−^ primary microglia were stained for Iba1. Compared with WT cells, *Clu*^−/−^ cells displayed more rounded cell bodies and shorter processes, resembling an activated or amoeboid morphology. Scale bars 20 μm. **(C)** Iba1 images were quantified, and results were normalized to WT and compared using Student’s *t*-test. Data are presented as the group mean ± SEM; *n* = 4/group. **(D)** WT and *Clu*^−/−^ primary microglia were treated with either 100 ng/mL LPS or vehicle alone for 24 hours and then compared for pro-IL-1β expression in the cell lysate. Results were normalized to the LPS-treated WT group and compared using one-way ANOVA with Tukey’s post hoc test. Data are presented as the group mean ± SEM; *n* = 4-5/group. **(E)** IMG cells were transfected with *Clu* siRNA or a scramble control, then treated with 100 ng/mL LPS or vehicle alone for 4 hours. Results were normalized to the LPS-treated Ctrl siRNA group and compared using one-way ANOVA with Tukey’s post hoc test. Data are presented as the group mean ± SEM; *n* = 5/group. (F-J) WT and *Clu*^−/−^ primary microglia were treated with 100 ng/mL LPS or vehicle alone for 24 hours, then analyzed by RT-qPCR for mRNA expression of microglia activation markers. Results were normalized to the LPS-treated WT group and compared using one-way ANOVA with Tukey’s post hoc test. Data are presented as the group mean ± SEM; *n* = 3-6/group. **(K-O)** IMG cells were transiently transfected with *Clu* siRNA or a scramble control. 48 hours post-transfection, cells were treated with 10 ng/mL LPS or vehicle alone for 1 hour, after which cells were analyzed for mRNA expression of microglia activation markers by RT-qPCR. Results were normalized to the LPS-treated control siRNA group and compared using one-way ANOVA with Tukey’s post hoc test. Data are presented as the group mean ± SEM; *n* = 3/group. \**p* < 0.05, \*\**p* < 0.01, \*\*\**p* < 0.001, \*\*\*\**p* < 0.0001.

To gain insight into the physiological role of microglia-derived CLU, primary microglia were cultured from both WT and *Clu*^−/−^ postnatal mice and compared for pro-IL-1β expression in cell lysate. The data revealed a greater increase in pro-IL-1β expression in *Clu*^−/−^ microglia than in WT microglia upon LPS treatment (Fig. 3D). In addition, *Clu* siRNA was transiently transfected into IMG cells to suppress CLU expression, with scrambled siRNA used as a control (Fig. S3). After 48 hours of transfection, IMG cells were treated with 10 ng/mL LPS or vehicle for 1 hour. Consistent with observations in primary microglia, loss of CLU was associated with elevated pro-IL-1β expression in cell lysates (Fig. 3E).

Furthermore, WT and *<u>Clu</u>*^−/−^ primary microglia were treated overnight with 100 ng/mL LPS or vehicle and compared for expression of proinflammatory cytokines and microglial activation markers using RT-qPCR. Compared with WT, LPS-treated *Clu*^−/−^ microglia exhibited a heightened inflammatory response, as evidenced by significantly elevated levels of the activation markers Iba1, CD11b, and TNF-α (Fig. 3F-J). Consistent with observations in primary microglia, loss of CLU was associated with an elevated inflammatory response in IMG cells, as indicated by significantly higher levels of Iba1, IL-1β, and IL-6 (Fig. 3K-O). Collectively, these findings demonstrate that loss of CLU intensified the LPS-induced inflammatory response in microglia and led to altered microglial morphology *in vitro*.

### Introduction of recombinant CLU (rCLU) increased microglial resistance to LPS stimulation

Based on the findings above, which indicate a strong association between CLU loss and elevated microglial inflammation, we hypothesized that introducing CLU could prevent microglial hyperactivation. To test this hypothesis, IMG cells were pre-incubated overnight with glycosylated, mature recombinant CLU (rCLU; Fig. 4A and Fig. S4) at a final concentration of 10 μg/mL or vehicle alone. Subsequently, cells were treated with 100 ng/mL LPS for 1 hour. Cell lysates were collected after treatment and analyzed for pro-IL-1β protein expression. A significant reduction in cytosolic pro-IL-1β levels was observed in IMG cells pre-incubated with rCLU compared with those pre-incubated with vehicle before LPS treatment (Fig. 4B). The cell culture medium was also analyzed for levels of two proinflammatory cytokines, IL-6 and TNF-α. IL-6 levels were significantly reduced with rCLU treatment (Fig. 4C); a similar trend was observed for TNF-α, though it did not reach statistical significance (Fig. 4D).

**Fig. 4.**
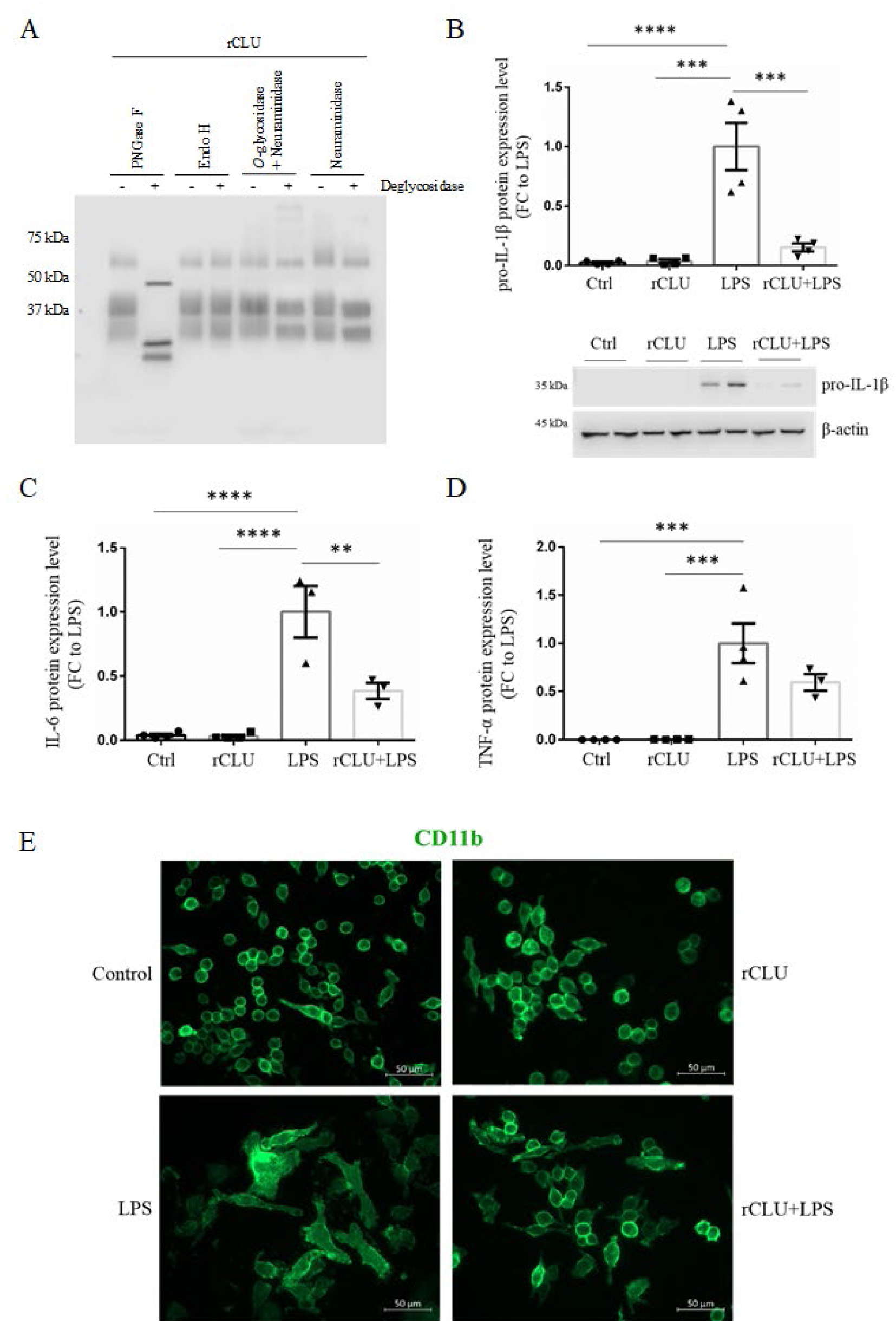
Recombinant CLU (rCLU) treatment attenuated microglial inflammatory responses to LPS stimulation. **(A)** Validation of rCLU expression and glycosylation profiles. PNGase F treatment confirmed that rCLU was highly glycosylated with N-linked glycans. Neuraminidase treatments confirmed that rCLU was sialylated. These PTM profiles are consistent with those of endogenous CLU expressed in mouse brains. **(B)** IMG cells were pretreated overnight with 10 μg/mL rCLU or vehicle alone, followed by LPS treatment for 1 hour. rCLU pretreatment significantly reduced pro-IL-1β expression in the cell lysate. Results were normalized to the LPS-treated group and compared using one-way ANOVA with Tukey’s post hoc test. Data are presented as the group mean ± SEM; *n* = 4/group. **(C-D)** IMG cells were pretreated overnight with 10 ng/mL rCLU or vehicle alone, then treated with LPS for 1 hour. Medium fractions were collected and analyzed for IL-6 and TNF-α expression levels using Bio-Plex cytokine assays. Results were normalized to the LPS-treated group and compared using one-way ANOVA with Tukey’s post hoc test. Data are presented as the group mean ± SEM; *n* = 4/group. **(E)** IMG cells were pretreated overnight with 10 μg/mL rCLU or vehicle alone, followed by LPS treatment for 4 hours and stained for CD11b for morphological comparisons. Vehicle alone or rCLU-treated control cells exhibited a narrow cell body; in contrast, LPS-treated IMG cells displayed significantly enlarged cell bodies. Cells pretreated with rCLU, followed by LPS treatment, showed a morphology resembling that of control cells. Scale bars 50 μm. \*\**p* < 0.01, \*\*\**p* < 0.001, \*\*\*\**p* < 0.0001.

In addition to biochemical analysis, morphological changes in microglia in response to LPS and the potential role of CLU in modulating these changes were also investigated. IMG cells were treated with 100 ng/mL LPS or vehicle for 4 hours, then stained with CD11b to assess morphology (Fig. 4E). LPS-treated cells acquired an amoeboid shape with an enlarged cell body compared with vehicle-treated cells. However, pretreatment of cells with 10 μg/mL rCLU overnight prevented microglia from undergoing LPS-induced morphological changes. In summary, as demonstrated by both biochemical and morphological observations, pretreatment with rCLU increased microglial resistance to LPS-mediated inflammation.

### Loss of CLU correlated with a significant reduction in Aβ plaque deposition in the cortex-hippocampal region of the *App*^NL-F/NL-F^ mouse model of AD

To investigate the influence of CLU on Aβ plaque deposition, sections from the left hemisphere were obtained from male and female *App*^NL-F/NL-F^ mice aged 10 and 15 months, with and without CLU. Campbell-Switzer staining revealed that at 10 months, plaque formation had just begun, with females exhibiting greater deposition than males (Fig. 5A and 5B). By 15 months, plaque deposition increased significantly, spreading throughout the cortex and select hippocampal regions, with consistently higher deposition in females. Compared with *App*^NL-F/NL-F^ x *Clu*^+/+^ mice, *App*^NL-F/NL-F^ x *Clu*^−/−^ mice exhibited a notable reduction in plaque deposition in both age groups, with females showing a more pronounced decrease than males (Fig. 5C and 5D). Remarkably, despite the overall reduction of plaques due to CLU loss, a distinct spatial pattern emerged, with *App*^NL-F/NL-F^ x *Clu*^−/−^mice displaying greater plaque accumulation in the subicular region.

**Fig. 5.**
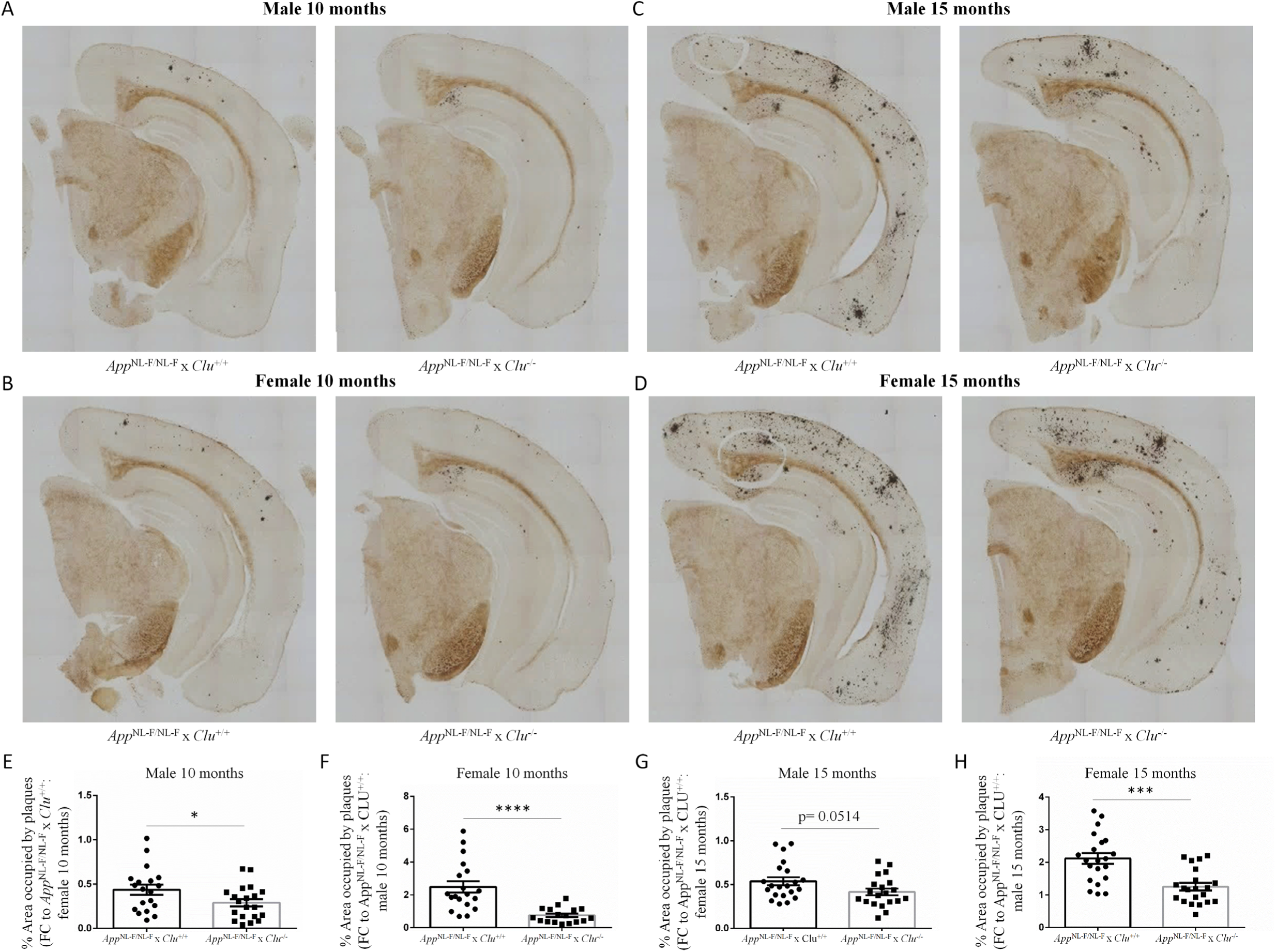
Loss of CLU reduced Aβ plaque deposition in *App*^NL-F/NL-F^ mice. **(A-D)** The left-brain hemispheres of 10- and 15-month-old male and female *App*^NL-F/NL-F^ × *Clu*^+/+^ and *App*^NL-F/NL-F^ × *Clu*^−/−^ mice were embedded together in a gelatin matrix, and the block was then coronally sectioned at 35 µm, encompassing the hippocampal region. Campbell-Switzer silver staining revealed plaque formation at 10 months and a significant increase by 15 months, with females showing more robust deposition than males. CLU loss was associated with reduced Aβ plaque deposition across all examined groups. **(E-H)** The percentage area of Aβ plaque coverage was compared between *App*^NL-F/NL-F^ × *Clu*^+/+^ and *App*^NL-F/NL-F^ × *Clu*^−/−^ mice for (E) male 10 months, (F) female 10 months, (G) male 15 months, and (H) female 15 months. Results were normalized to brains of the specific strain/sex/age on the same section as indicated on the y-axis, allowing comparisons of plaque deposition across animals at the same anatomical locations. Results were then analyzed via Student’s *t*-test. Data are presented as the group mean ± SEM; *n* = 21 sections/group. \**p* < 0.05, \*\**p* < 0.01, \*\*\**p* < 0.001, \*\*\*\**p* < 0.0001.

To account for anatomical variations in Aβ plaque coverage across regions, group comparisons were performed by normalizing to the sections on the same blocks. The analysis showed that in *App*^NL-F/NL-F^ mice, CLU loss led to a significant reduction in Aβ plaque formation in both male (Fig. 5E) and female mice (Fig. 5F) at 10 months of age. Furthermore, a notable reduction was observed in 15-month-old mice, reaching statistical significance only in females (Fig. 5G), and showing a borderline *p*-value in males (Fig. 5H). These findings indicate a strong positive correlation between CLU loss and reduced Aβ plaque deposition, with a more pronounced reduction in females than in males.

### Loss of CLU led to a notable decline in learning and memory activities in the *App*^NL-F/NL-F^ mouse model of AD

To assess the impact of CLU on cognitive functions, a two-trial Y-maze test was performed. Mice explored the Y-maze with one arm blocked for 5 minutes, then returned to their cage. After 30 minutes, they revisited the maze and freely explored all three arms for 5 minutes (Fig. 6A). A decrease in time spent exploring the novel arm suggests diminished spatial learning capacity and recognition memory (*30*). A trend, although not statistically significant (*p* = 0.0558), was observed, with *App*^NL-F/NL-F^ x *Clu*^−/−^ mice spending less time in the novel arm than *App*^NL-F/NL-F^ x *Clu*^+/+^ mice (Fig. 6B). Additionally, no significant differences were noted in the total distance traveled within the maze between the two genotypes, indicating comparable mobility (Fig. 6C).

**Fig. 6.**
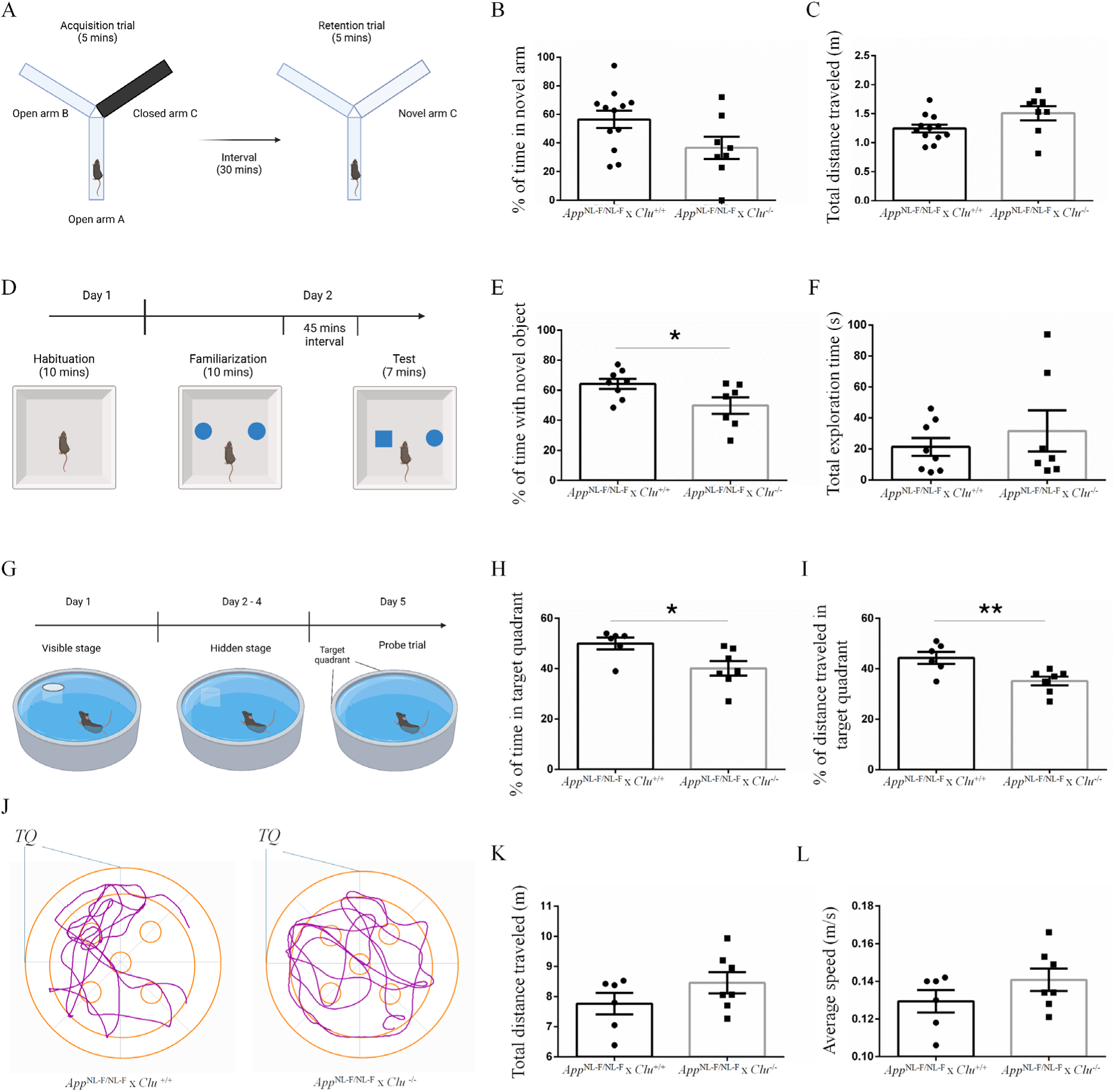
Loss of CLU impaired learning and memory activities in *App*^NL-F/NL-F^ mice. **(A)** Schematic of the Y-maze two-trial behavioral test paradigm. **(B-C)** The total distance traveled in the maze was similar across groups; however, *App*^NL-F/NL-F^ × *Clu*^−/−^ mice showed a non-significant trend toward spending less time in the novel arm than *App*^NL-F/NL-F^ × *Clu*^+/+^ mice. **(D)** Schematic of the NOR behavioral test paradigm. **(E-F)** Total exploration time was similar between the groups; however, *App*^NL-F/NL-F^ × *Clu*^−/−^ mice spent significantly less time exploring novel objects than *App*^NL-F/NL-F^ × *Clu*^+/+^ mice. Data are presented as the group mean ± SEM; *n* = 8-12 mice/group. **(G)** Schematic of the MWM behavioral test paradigm. **(H-I)** Total distance traveled and average speed did not differ significantly between the groups. **(J-L)** *App*^NL-F/NL-F^ × *Clu*^−/−^ mice spent significantly less time and traveled a shorter distance in the target quadrant (TQ) than *App*^NL-F/NL-F^ × *Clu*^+/+^ mice. Data are presented as the group mean ± SEM; *n* = 6-7 mice/group. **J)** Track plots illustrate preference for the TQ in *App*^NL-F/NL-F^ × *Clu*^+/+^ but not in *App*^NL-F/NL-F^ × *Clu*^−/−^ mice. Schematic figures were created using BioRender. Results were analyzed via Student’s *t*-test. \**p* < 0.05, \*\**p* < 0.01.

Next, recognition memory was assessed using the novel object recognition (NOR) test. Mice were first habituated in an empty open field for 10 minutes. After 24 hours, two identical objects were placed 5 cm from the wall. Each mouse was then placed in the field for 10 minutes. After a 45-minute break, one object was replaced with a novel object. Mice had 7 minutes to explore, and interaction time with both objects was recorded (Fig. 6D). A preference for a novel object over a familiar one reflects the ability to remember and recall previously encountered objects (*31*). Compared with *App*^NL-F/NL-F^ x *Clu*^+/+^ mice, *App*^NL-F/NL-F^ x *Clu*^−/−^ spent significantly less time exploring the novel object, indicating impaired recognition memory with loss of CLU (Fig. 6E). Furthermore, total exploration time was comparable between the two groups, ruling out nonspecific differences due to preferential bias (Fig. 6F).

Finally, the impact of CLU on spatial learning and long-term memory was assessed using the Morris water maze (MWM) test. Training comprised two stages: Day 1 (visible stage - 5 trials/mouse) with the platform above clear water, positioned at different locations; Days 2-4 (hidden training - 5 trials/mouse) with the platform in a fixed position (target quadrant), submerged in opaque water. On Day 5 (probe trial - 1 trial/mouse), the platform was removed, and mice started from a designated quadrant (Fig. 6G). Less time and distance in the target quadrant indicate impaired cognition (*32*). *App*^NL-F/NL-F^ x *Clu*^−/−^ mice spent significantly less time (Fig. 6H) and covered a shorter distance (Fig. 6I) in the target quadrant during the probe trial than *App*^NL-F/NL-F^ x *Clu*^+/+^ mice. Tracking plots consistently demonstrate a preference for the target quadrant in *App*^NL-F/NL-F^ x *Clu*^+/+^ mice but not in *App*^NL-F/NL-F^ x *Clu*^−/−^ mice (Fig. 6J). The total distance traveled (Fig. 6K) and average speed (Fig. 6L) were similar between the two groups, indicating comparable mobility. Collectively, these behavioral outcomes strongly support that CLU deficiency can lead to reduced cognitive function in *App*^NL-F/NL-F^ mice.

### Loss of CLU increased microglial activation and inflammation in both WT and the *App*^NL-F/NL-F^ mouse model of AD

Given the substantial *in vitro* evidence of heightened microglial activation and inflammation resulting from CLU deficiency (Fig. 3), we sought to determine whether a similar phenotype is observed *in vivo*. To this end, cortical tissues were isolated from both male and female 15-month-old WT and *Clu*^−/−^ mice. CD11b, a key microglial surface protein whose expression correlates with the intensity of microglial activation (*33*), was examined by Western blotting, which revealed significantly higher CD11b expression in *Clu*^−/−^ mice than in WT mice (Fig. 7A). In addition to surface marker alterations, activated microglia release substantial amounts of proinflammatory cytokines, including IL-1β, IL-6, and TNF-α (*34*). To compare cytokine production between WT and *Clu*^−/−^ mice, RT-qPCR was performed to measure mRNA levels of the two most commonly reported cytokines, IL-1β and TNF-α. A significant increase in expression of these cytokines was observed in *Clu*^−/−^ mice (Fig. 7B and 7C). These findings indicate a certain level of microglial inflammation in *Clu*^−/−^ mice compared with WT mice.

**Fig. 7.**
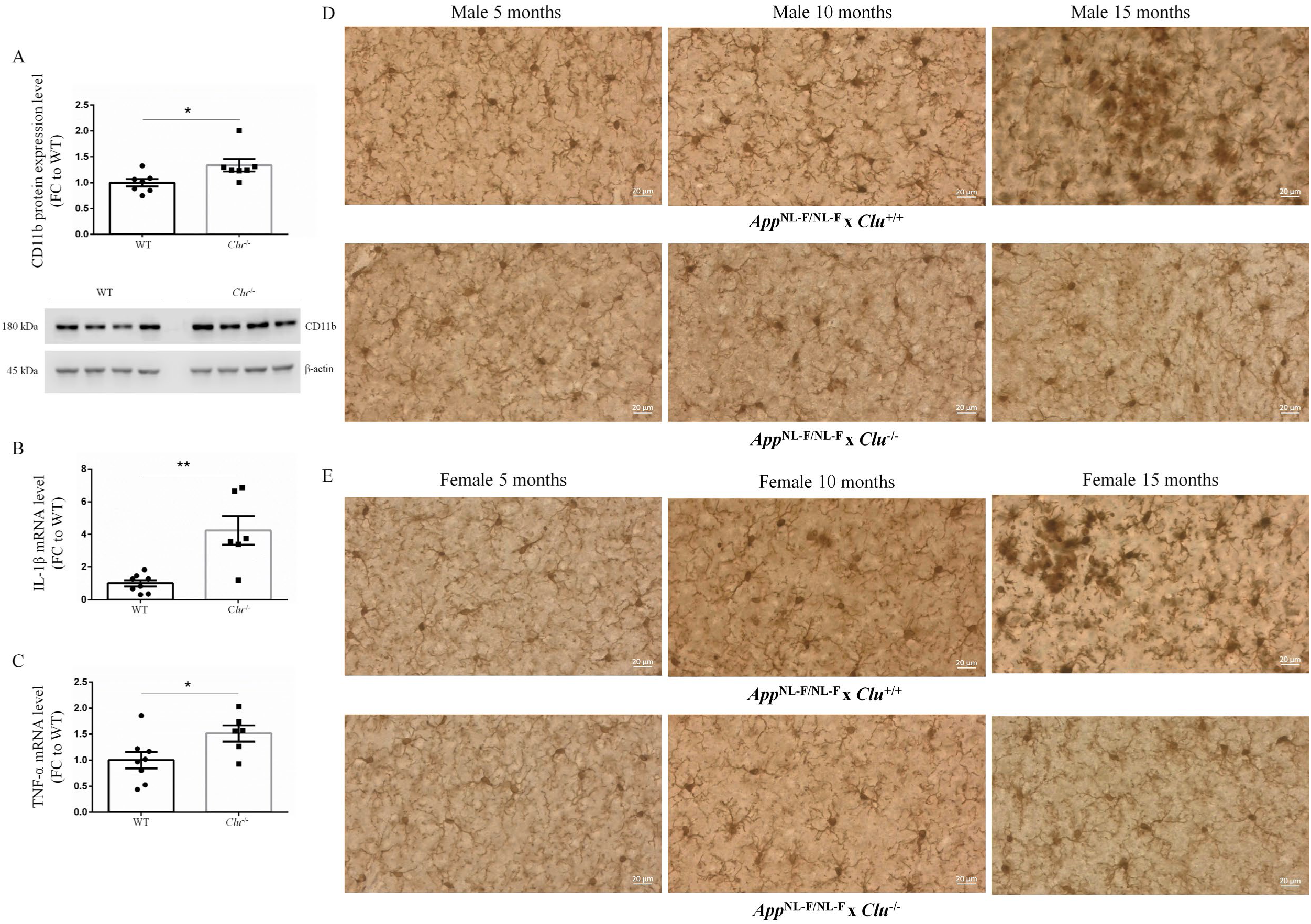
Loss of CLU promoted microglia activation and inflammation in both WT and *App*^NL-F/NL-F^ mice. **(A-C)** WT and Clu^−/−^ mice cortical tissue lysates were compared for (A) CD11b protein expression levels, (B) IL-1β mRNA levels, and (C) TNF-α mRNA levels. Results were normalized to WT and compared using Student’s *t*-test. Data are presented as the group mean ± SEM; *n* = 6-8 mice/group. \**p* < 0.05, \*\**p* < 0.01. **(D-E)** Longitudinal morphological phenotypes of microglia in *App*^NL-F/NL-F^ × *Clu*^+/+^ and *App*^NL-F/NL-F^ × *Clu*^−/−^ mice at the age of 5 months, 10 months, or 15 months in (D) males and (E) females. In *App*^NL-F/NL-F^ × *Clu*^+/+^ mice, microglia transitioned from ramified to reactive and amoeboid morphologies over time, whereas *App*^NL-F/NL-F^ × *Clu*^−/−^ mice did not display these dynamic changes.

A longitudinal analysis of microglial morphology was conducted in *App*^NL-F/NL-F^ mice with and without CLU expression. In *App*^NL-F/NL-F^ x *Clu*^+/+^ mice, microglia exhibited a predominantly ramified morphology at 5 months of age, followed by a progressive increase in reactive and amoeboid phenotypes over time, indicative of heightened microglial activation and inflammation with disease progression. In contrast, *App*^NL-F/NL-F^ x *Clu*^−/−^ mice failed to show this morphological transition and instead displayed an early and sustained presence of reactive microglia (Fig. 7D-E). These findings suggest that the absence of CLU promotes an early shift toward a pro-inflammatory microglial state, which may lead to chronic activation and eventual functional desensitization, as evidenced by a diminished morphological response to Aβ plaque deposition in later stages.

### Loss of CLU disrupted the balance between excitatory and inhibitory neurons in both WT and the *App*^NL-F/NL-F^ mouse model of AD

Despite significantly reducing plaque deposition, CLU loss in *App*^NL-F/NL-F^ mice led to a decline in cognitive performance. This suggests a cognitive regulatory pathway influenced by CLU, distinct from mechanisms mediated by Aβ pathology. To elucidate such pathways, snRNA-seq analysis was performed on brain cortical samples from 15-month-old male mice. First, a comparison was made between WT and *Clu*^−/−^ mice. t-Distributed Stochastic Neighbor Embedding (t-SNE) plots were generated, with each dot representing an individual cell. Clusters were color-coded to represent distinct cell types, differentiated by the similarity of their gene expression profiles. Eight distinct clusters were identified, representing excitatory neurons, inhibitory neurons, astrocytes, microglia, oligodendrocytes, oligodendrocyte precursor cells, and endothelial cells, with one cluster remaining unclassified (Fig. 8A). When the tSNE plots were color coded to represent cells from WT cortex in red and cells from *Clu*^−/−^ cortex in green, clear differences in the relative numbers of excitatory and inhibitory neurons between the two genotypes became apparent (Fig. 8B). To further illustrate these findings, pie charts were used to present the percentage of each cell type relative to the total counts. When compared to WT cortex, *Clu*^−/−^ cortex showed a significant decrease in the proportion of excitatory neurons (ExN) from 53% to 35%, with a simultaneous increase in the proportion of inhibitory neurons (InN) from 10% to 21% (Fig. 8C and 8D). In addition, the percentages of both oligodendrocytes (Oligo) and oligodendrocyte precursor cells (OPC) increased in *Clu*^−/−^ cortex compared with WT cortex. KEGG enrichment analysis of DEGs revealed that the top three biological pathways affected by CLU deficiency were axon guidance, Ca^2+^ signaling pathway, and long-term potentiation (Fig. 8E).

**Fig. 8.**
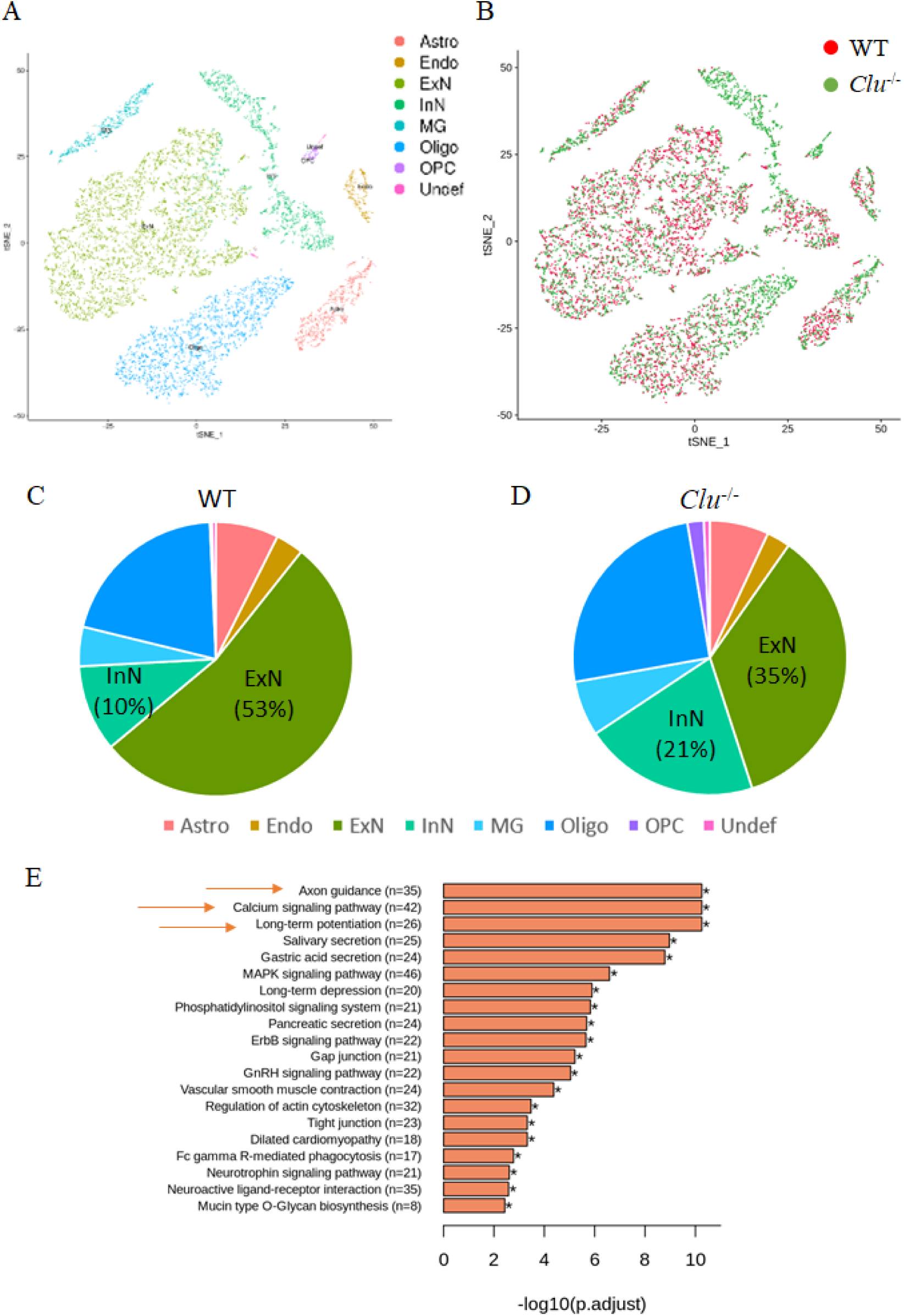
snRNA-seq data revealed a disruption in the balance between excitatory and inhibitory neurons in *Clu*^−/−^ mice compared to WT mice. **(A)** tSNE plots were generated from the brains of WT and *Clu*^−/−^ mice. Each dot represents a single cell. Cells are color-coded, with each color representing a distinct cell type. A total of eight cell populations were identified, with one cluster unidentified. **(B)** tSNE plots show cortical cell clusters from WT and *Clu*^−/−^ mice, with each genotype assigned to a distinct color. **(C-D)** Pie charts show the relative proportions of cell types in WT and *Clu*^−/−^ mice, with percentages of excitatory (ExN) and inhibitory (InN) neurons indicated. **(E)** KEGG enrichment analysis of DEGs, with the top three biological pathways highlighted by orange arrows.

Subsequently, similar comparisons were made between the brain cortices of *App*^NL-F/NL-F^ x *CLU*^+/+^ and *App*^NL-F/NL-F^ x *Clu*^−/−^ mice. t-SNE plots were generated and color-coded to represent distinct cell clusters. Eight clusters were identified, with one remaining unclassified (Fig. 9A). As in the *Clu*^−/−^ vs WT comparison, a clear disparity in the relative numbers of excitatory and inhibitory neurons was found between *App*^NL-F/NL-F^ x *Clu*^+/+^ cortex and *App*^NL-F/NL-F^ x *Clu*^−/−^ cortex (Fig. 9B). A significant decrease in the proportion of ExN, from 55% to 24%, and an increase in the proportion of InN, from 10% to 24%, were observed in *App*^NL-F/NL-F^ x *Clu*^−/−^ cortex compared with *App*^NL-F/NL-F^ x *Clu*^+/+^ cortex (Fig. 9C and 9D). The greater decrease in ExN in *App*^NL-F/NL-F^ x *Clu*^−/−^ vs *App*^NL-F/NL-F^ x *Clu*^+/+^ cortex, compared with the decrease in *Clu*^−/−^ vs WT cortex, indicates an amplified impact of CLU loss in the context of AD. Moreover, consistent with the *Clu*^−/−^ and WT comparison, the proportions of Oligo and OPC increased in *App*^NL-F/NL-F^ x *Clu*^−/−^ cortex compared with *App*^NL-F/NL-F^ x *Clu*^+/+^ cortex. KEGG enrichment analysis of DEGs revealed that the top three biological pathways affected by CLU deficiency in *App*^NL-F/NL-F^ mice are identical to those observed in *Clu*^−/−^vs WT comparisons, underscoring the crucial role of CLU in promoting neuronal homeostasis and cognitive function (Fig. 9E).

**Fig. 9.**
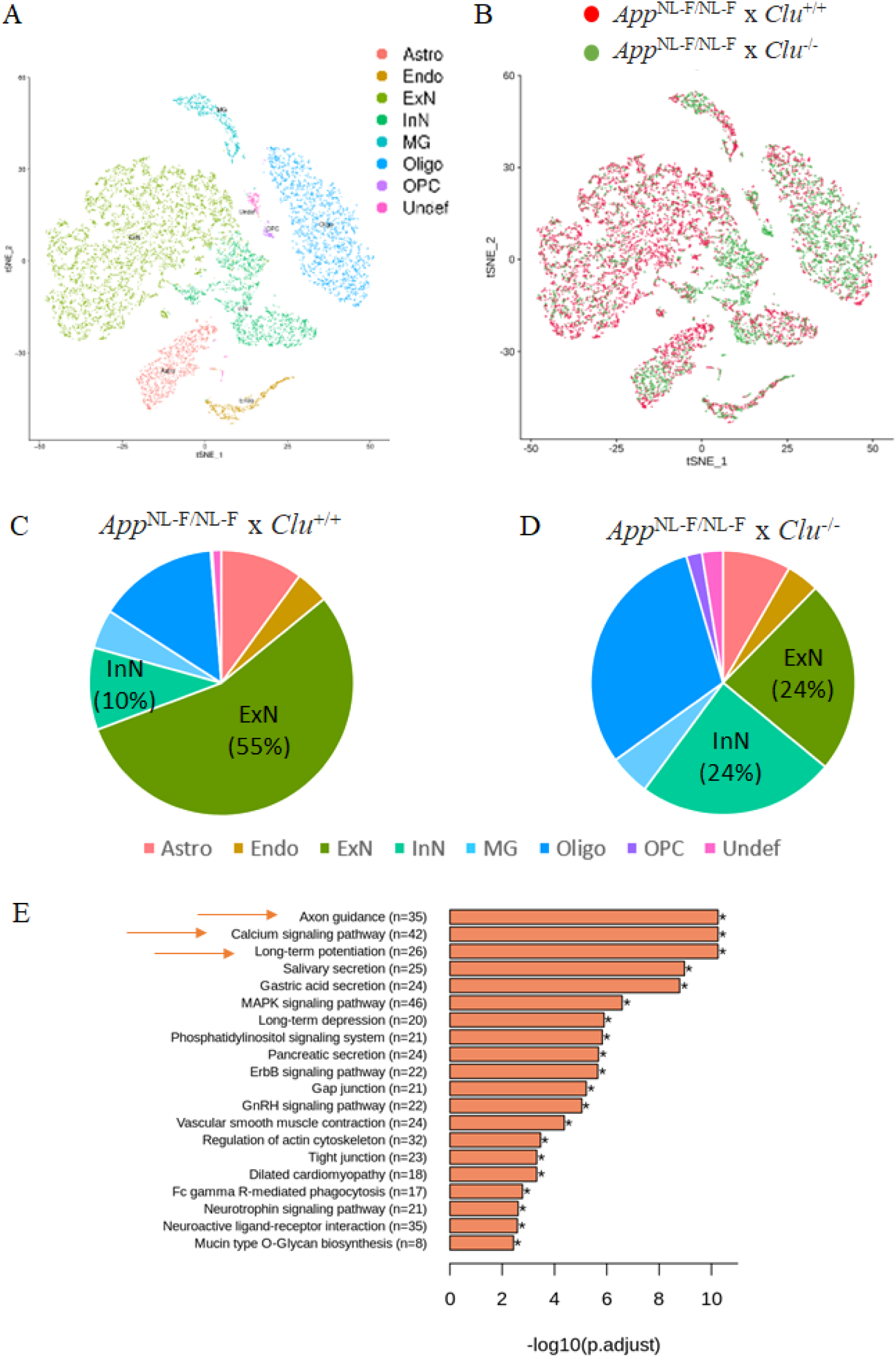
snRNA-seq data revealed a disruption in the balance between excitatory and inhibitory neurons due to CLU loss in *App*^NL-F/NL-F^ mice. **(A)** tSNE plots were generated from the brains of *App*^NL-F/NL-F^ × *Clu*^+/+^ and *App*^NL-F/NL-F^ × *Clu*^−/−^ mice. Each dot represents a single cell. Cells are color-coded, with each color representing a distinct cell type. A total of eight cell populations were identified, with one cluster unidentified. **(B)** tSNE plots were color-coded to represent different strains. **(C-D)** Pie charts show the relative proportions of cell populations in *App*^NL-F/NL-F^ × *Clu*^+/+^ and *App*^NL-F/NL-F^ × *Clu*^−/−^ mice, with percentages of ExN and InN neurons indicated. **(E)** KEGG enrichment analysis of DEGs demonstrates the altered biological pathways; orange arrows highlight the top three pathways.

## DISCUSSION

Astrocytes are well-established as the primary source of CLU in the brain (*35*). However, microglial CLU expression remains poorly characterized. Given the existence of multiple CLU isoforms, we conducted a detailed analysis of microglial *Clu* mRNA and protein expression under basal and LPS-induced inflammatory conditions. CLU expression in resting microglia was minimal but substantially increased upon LPS stimulation in a dose- and time-dependent manner. The detected bands correspond to the high-mannose intermediate (~60 kDa), mCLUα (~34–37 kDa), and mCLUβ (~36–39 kDa).

To assess the functional relevance of LPS-induced CLU upregulation, we analyzed primary microglia from WT and *Clu*^−/−^ mice, and IMG cells transiently transfected with *Clu* or scrambled siRNA. In both models, CLU deficiency heightened inflammatory responses, indicating that microglial CLU acts as an anti-inflammatory modulator. This supports previous findings that peripheral CLU from runner plasma dampens brain inflammation via vascular interactions (*36*). Mechanistically, increased microglial activation in the absence of CLU, compared with WT microglia, led us to hypothesize that CLU functions as a negative feedback inhibitory regulator of microglial activation and promotes microglial resolution. Consistent with this hypothesis, recombinant CLU supplementation significantly downregulated microglial activation and inflammation, as evidenced by morphological changes and inflammatory cytokine levels. In contrast, a previous study using human CLU on rat microglia reported increased activation (*37*), likely due to cross-species reactivity. The use of species-matched rCLU in the present study provides physiologically relevant evidence supporting an anti-inflammatory role for CLU in microglia.

The relationship between CLU and Aβ pathology is complex. Earlier studies documented that CLU colocalizes with Aβ plaques and acts as a molecular chaperone to prevent toxic aggregation (*38–41*). However, recent literature presents a mixed picture, with some studies reporting CLU overexpression and reduced plaque burden (*42*), while others report increased Aβ deposition with CLU (*43, 44*). These conflicting outcomes likely reflect differences in experimental models (*4*). To further clarify the potential role of CLU in Aβ pathology, we analyzed the impact of CLU on Aβ plaque formation and accumulation in an *App* knock-in (*App*^NL-F/NL-F^) mouse model of familial AD (*45*). This model avoids APP overexpression and more accurately recapitulates clinical AD pathology by producing age-dependent Aβ accumulation without generating excess APP fragments (*46*). Loss of CLU in *App*^NL-F/NL-F^ mice significantly reduced overall brain Aβ plaque deposition, with a greater effect in females. Regionally, CLU deficiency reduced cortical plaques but led to persistent accumulation in the subiculum. This pattern may stem from regional differences in vascular clearance capacity, with the hippocampus exhibiting lower baseline perfusion, oxygenation, and neurovascular coupling than the cortex, which may limit Aβ clearance in subicular regions (*47*).

While the role of CLU in Aβ deposition is controversial, a prevailing consensus in the literature indicates that reducing Aβ levels improves cognitive abilities across various AD animal models (*48–55*). However, the findings from the current study diverge from this consensus. Despite a significant reduction in both diffuse and amyloid core plaques in CLU-deficient mice, these mice exhibited markedly worse performance on various cognitive tests, suggesting that Aβ plaque-independent cognitive mechanisms are involved. This aligns with clinical efforts targeting Aβ plaques in AD patients, in which significant reductions in brain Aβ deposits failed to translate into cognitive benefits (*56*). One hypothesis proposed to explain Aβ-based therapeutic failures in improving cognition is that treatment may have been initiated too late in the disease pathophysiology; targeting the preclinical phase of the disease, characterized by early Aβ deposition, may be more likely to have a positive effect on cognitive symptoms (*57, 58*). However, the findings from our study, involving 10-month-old AD mice just beginning to show plaque formation, consistently demonstrate no cognitive benefit from reducing plaque deposition associated with CLU loss. This suggests the existence of alternative Aβ plaque-independent mechanisms by which CLU may affect cognitive function.

To investigate how CLU regulates cognition, snRNA-seq was performed. Strikingly, CLU deficiency was associated with a significant disruption of the balance between excitatory and inhibitory neurons and a notable reduction in the number of excitatory neurons. These observations of neuronal dyshomeostasis resulting from CLU deficiency were even more pronounced in *App*^NL-F/NL-F^ mice. Excitatory neurons have been found to be more vulnerable and preferentially impaired in AD (*59, 60*). The specific loss of excitatory neurons and its correlation with memory deficits have been reported in both Aβ-based (*61*) and tau-based AD mouse models (*62*). Therefore, the significant reduction in excitatory neurons could have contributed to the cognitive impairments observed in CLU deficiency. KEGG enrichment analysis of DEGs further revealed that the top three biological pathways altered by CLU deficiency were axon guidance, Ca^2+^ signaling pathway, and long-term potentiation, providing additional support for a crucial role of CLU in maintaining neuronal homeostasis and, consequently, learning and memory function. It remains to be determined whether the selective loss of excitatory neurons was caused, at least in part, by chronic inflammation resulting from CLU deficiency. This hypothesis is strongly supported by recent studies showing that excitatory neurons are more vulnerable to inflammatory stress and neurodegeneration than inhibitory neurons. This heightened sensitivity could be linked to preferential disruptions of mitochondrial and synaptic MAPK signaling in excitatory neurons (*63*). Other mechanisms may also be involved. Support for this possibility comes from a recent study examining the role of CLU in synaptic transmission. *Ex vivo* recordings of miniature excitatory and inhibitory postsynaptic currents (mEPSCs and mIPSCs) were conducted on hippocampal granule neurons from postnatal day 28–30 *Clu*^−/−^ mice and their WT littermate controls. A significant decrease in mEPSC frequency was observed in *Clu*^−/−^ neurons, whereas mIPSC frequency and amplitude remained unaffected (*42*).

Finally, another notable finding was that CLU deficiency was associated with significant increases in the number of oligodendrocytes and oligodendrocyte precursor cells (OPCs). OPCs differentiate into myelinating oligodendrocytes (*64*) and have been shown to proliferate in response to neuronal injury, facilitating remyelination (*65*). The cause of this increase resulting from CLU deficiency was not investigated in this study; however, similar phenotypes have been frequently observed in aging and AD brains, likely reflecting a compensatory attempt at repair in response to neuronal degeneration and death, despite ultimately being dysfunctional (*66–68*). Moreover, recent studies have indicated a novel role of OPCs as dynamic phagocytes involved in myelin debris clearance and Aβ phagocytosis (*69–71*). Furthermore, although unrelated to Aβ, the ability of mature oligodendrocytes to phagocytose carbon particles was reported as early as 1989, yet this phenomenon remained largely unexplored in the years that followed (*72*). Therefore, it is conceivable that the proliferation of oligodendrocytes and OPCs resulting from CLU deficiency could enhance Aβ phagocytosis, which may underlie the reduction in Aβ plaque deposition observed in the *App*^NL-F/NL-F^ x *Clu*^−/−^ mice.

In summary, this study reveals novel roles for CLU in maintaining brain health and function. Microglia expressed low levels of CLU under resting conditions; however, microglial CLU expression and secretion significantly increased in response to pro-inflammatory stimulation. These observations, together with the finding that CLU deficiency increased microglial activation and inflammation, indicate that CLU functions as a negative feedback inhibitor of microglial activity. Moreover, CLU deficiency was found to disrupt the balance between excitatory and inhibitory neurons, largely due to the preferential loss of excitatory neurons, which was accompanied by increased oligodendrogenesis and OPC proliferation. Furthermore, CLU deficiency was found to reduce Aβ plaque accumulation while impairing cognitive function in AD mouse models. Future studies to clarify how these phenotypes are interconnected (Fig. 10) will shed important light not only on the AD risk mechanism associated with CLU polymorphisms but also on the mechanisms of AD pathogenesis in general.

**Fig. 10.**
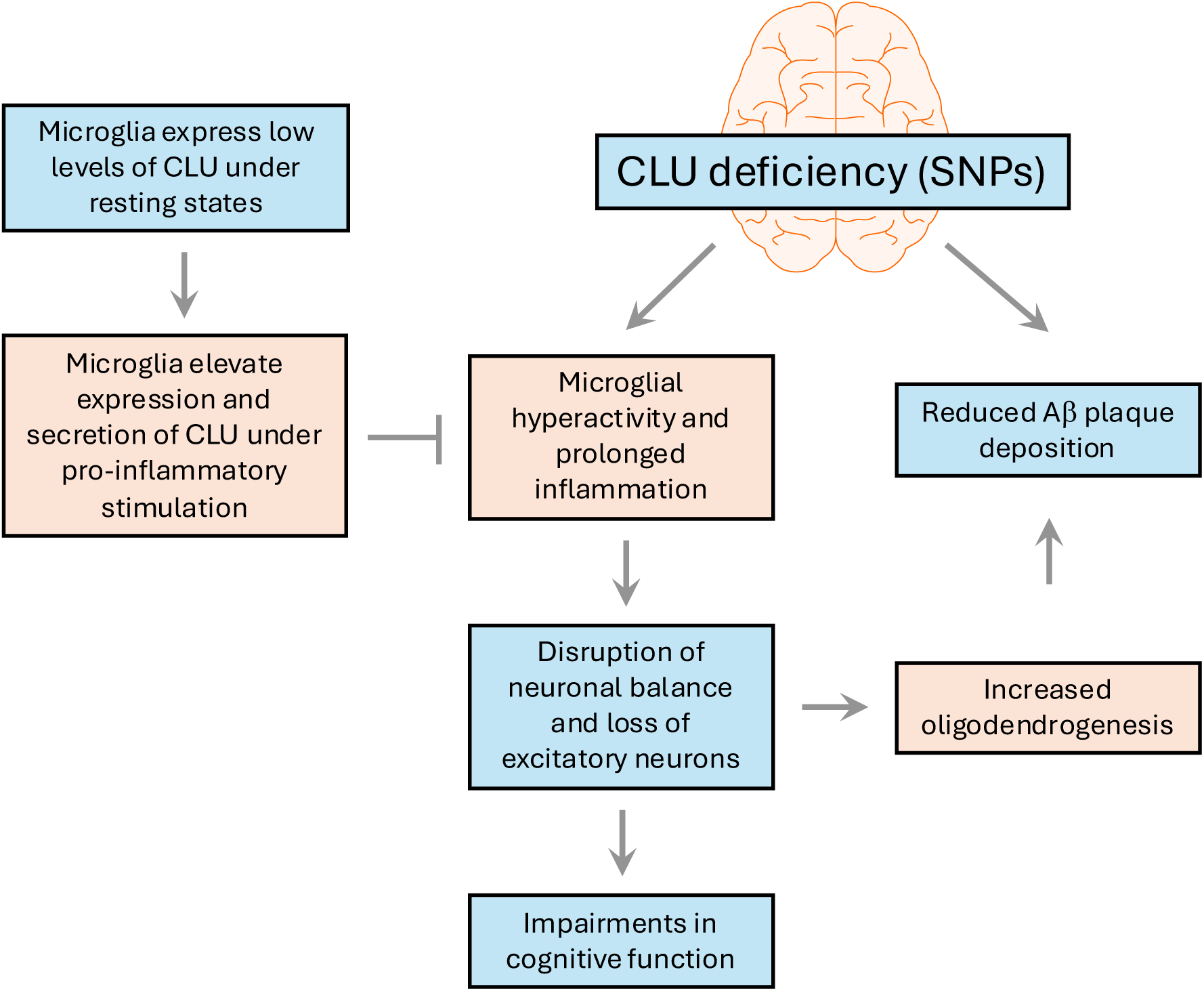
Summary of findings of the present study and mechanistic hypothesis. Microglia express low levels of CLU under resting conditions. When stimulated by pro-inflammatory signals, microglia increase CLU expression and secretion, and the resulting rise in CLU serves as a negative feedback inhibitor of microglial activation and inflammation. CLU deficiency associated with AD-risk SNPs can lead to excessive microglial activity and prolonged inflammation, which can further disrupt the balance between excitatory and inhibitory neurons, largely due to the loss of the most vulnerable excitatory neurons. Neuronal degeneration and death stimulate oligodendrogenesis and proliferation of oligodendrocyte precursor cells (OPCs) as part of a compensatory repair response. Increases in oligodendrocytes and OPCs can further enhance phagocytosis of Aβ plaques, thereby reducing plaque deposition. Disruption of neuronal homeostasis and excitatory neuronal death ultimately lead to impairments in cognitive function associated with AD.

## MATERIALS AND METHODS

### Animals

All animal procedures were approved by the Institutional Animal Care and Use Committee at the University of Kansas (Animal Use Statement: 220-04) and were performed in accordance with the National Institutes of Health (NIH) Guide for the Care and Use of Laboratory Animals. Mice were housed in a temperature-controlled environment with a 12-hour light/dark cycle and provided *ad libitum* access to food and water. The current study used three mutant mouse models, which are summarized below.

#### Generation of novel CLU^−/−^ mouse model

Our previous study demonstrated that while the commercially available *Clu*^−/−^ mouse model (JAX stock #005642), which was originally described by McLaughlin (*73*), lacks the mRNA transcript that produces the major mature isoform of CLU, several ‘minor’ isoforms were still detected in this model (*25*). To address these issues and generate a mouse model with complete loss of CLU function, lacking all isoforms, a novel complete protein null *Clu* knockout mouse model was generated using CRISPR technology (Fig. S5). CRISPR reagents were designed and validated (*74*), which targeted sequences in the *Clu* locus, flanking all coding exons: intron 1 (CRISPR target sequence TTGGCAGAGTATGCTAGTTCAGG; mm39 chr14:66,208,357-66,208,379); and the 3’ untranslated region of exon 9 (CRISPR target sequence AAGGGACACGTATCGCAAGGCGG; mm39 chr14:66,218,980-66,219,002). These CRISPR reagents were co-electroporated or co-injected as ribonucleoprotein complexes (Integrated DNA Technologies) into C57BL/6J zygotes as described (*75*). Founder mice with the deletion between the CRISPR targets, generated via non-homologous end joining and spanning 10,628 base pairs between these two target sites, were identified via PCR and sequencing (Fig. S5). One founder mouse was used to expand the deletion allele for subsequent matings. All the data involving the use of *Clu*^−/−^ primary microglia culture were derived from this new model. Except for the data presented in Fig 7A-C, which used the JAX model, all other data generated in *Clu*^−/−^ primary microglia culture and *App*^NL-F/NL-F^ × *Clu*^−/−^ double mutant mice were derived from this newly generated model.

#### App^NL-F/NL-F^ knock-in mouse model

The *App*^NL-F/NL-F^ knock-in mouse model was obtained from the RIKEN BioResource Research Center (BRC No. RBRC06343). As illustrated in Fig. S6, the mouse Aβ sequence was humanized by replacing three amino acids, followed by the introduction of Swedish and Beyreuther/Iberian mutations using knock-in technology. Genotyping was performed with minor modifications to the PCR cycling conditions from the supplier’s protocol to better distinguish between WT and *App*^NL-F/NL-F^ mice.

#### App^NL-F/NL-F^ × CLU^−/−^ double mutant mouse model

The *App*^NL-F/NL-F^ × *Clu*^−/−^ double mutant mouse model was generated by crossing *App*^NL-F/NL-F^ mice with *Clu*^+/−^ mice. *App*^NL-F/NL-F^ × *Clu*^+/+^ mice generated during breeding were used as controls for *App*^NL-F/NL-F^ × *Clu*^−/−^ mice in studies evaluating the impact of CLU loss on cognitive function and the development of AD-like pathology in the *App*^NL-F/NL-F^ mouse model of AD.

### Primary cultures of mouse microglia and astrocytes

Primary cultures of mouse microglia and astrocytes were prepared from postnatal pups of WT or *Clu*^−/−^ mice on day 0-2. Cortical tissues from mouse brains were dissected and collected in Hank’s balanced salt solution (HBSS) (Gibco, Cat #14170112). The tissues were then treated with 0.05% trypsin (Gibco, Cat #25300062) for 5 minutes at 37 °C. To neutralize the trypsin, astrocyte medium composed of Dulbecco’s modified Eagle’s medium (DMEM) (Gibco, Cat #11965092) supplemented with 10% fetal bovine serum (FBS) (Gibco, Cat #10437-028) and 1% penicillin/streptomycin (P/S) (Gibco, Cat #15070063) was added. The tissues were dissociated by repeatedly passing them through a series of fire-polished constricted Pasteur pipettes. The cortical cells were then plated onto poly-D-lysine (PDL) (Sigma, Cat #P7280) coated T-75 flasks at a density of 1 × 10^6^ cells/mL and incubated at 37 °C in a humidified 5% CO_2_ atmosphere. The medium was changed the following day, and the cells were left undisturbed for a total of 10 days, when microglia visibly grew on top of the astrocytic layer. On the 10^th^ day, the cell cultures were shaken at 200 rpm for 2 hours to remove microglia. The supernatant, containing the microglia cells, was collected and centrifuged at 170 x g for 10 minutes. The pellet was then resuspended in microglia medium (ScienCell, Cat #1901) and plated. For biochemical experiments, microglia cells were plated at a density of 5 × 10^5^ cells/mL, while for imaging experiments, they were plated at a density of 2 × 10^5^ cells/mL on PDL-coated cell culture dishes. Plated microglia cells were ready for experiments after 5 days. To prepare astrocytes, the T-75 flasks were further shaken overnight at 200 rpm to detach oligodendrocytes. The next day, the astrocytes were trypsinized and plated at a similar density to that mentioned above on PDL-coated plates. The astrocytes were maintained in astrocyte medium and were ready for use after 5 days.

### Mouse microglial cell line

IMG cell line was obtained from Sigma Aldrich (Cat #SCC134). IMG cells were cultured in high-glucose DMEM (Sigma, Cat #D6546), supplemented with 10% FBS, 1% L-glutamine, and 1% P/S. The cells were maintained at 37°C in a humidified 5% CO_2_ atmosphere. Cells with a passage number below 60 and in the exponential growth phase were utilized in the current study.

### siRNA transfection in IMG cells

*Clu* siRNA (Santa Cruz, Cat #sc-43689) and scramble control siRNA (Santa Cruz, Cat #sc-37007) were transfected into IMG cells using jetPRIME transfection reagent (jetPRIME, Cat #101000046) following the manufacturer’s instructions. Briefly, 10 nM *Clu* or control siRNA was combined with the transfection reagent and buffer, incubated at room temperature (RT) for 10 minutes, and added dropwise to the cell culture. After 48 hours of transfection, the cells were treated with 100 ng/mL LPS or vehicle for 4 hours for Western blot analysis and 10 ng/mL LPS for 1 hour for gene expression analysis.

### RT-qPCR gene expression

Total RNA was extracted from cell cultures or cortical tissue using the PureLink RNA Mini Kit (Thermo, Cat #12183018A). For cell cultures, the cells were washed twice with phosphate-buffered saline (PBS) (pH 7.4, Thermo Fisher Scientific, Cat #10010), homogenized with 400 μL TRIzol Reagent (Thermo, Cat #15596026), and collected in a sterile tube. To each tube, 80 μL chloroform was added, mixed by inversion, and centrifuged. For cortical tissue, the tissue was homogenized with 1 scoop of pre-chilled 400 μm Silica Beads (OPS Diagnostics, Cat #BMBG 400-200-05) and 200 μL TRIzol Reagent in a pre-chilled Bullet blender at speed 6 for 1 minute. An additional 400 μL TRIzol Reagent was added, and the homogenate was re-homogenized for 1 minute at speed 2. The homogenate was then centrifuged at 10,000 × g for 1 minute at 4 °C. The resulting supernatant was transferred to a new microcentrifuge tube, and 500 μL was collected. To each sample, 100 μL chloroform was added, mixed by inversion for 15 seconds, and centrifuged. After the chloroform mixing step, the downstream process was the same for both cell and tissue samples. The samples were centrifuged at 12,000 × g for 15 minutes at 4 °C, and approximately 100 μL and 200 μL of the upper supernatant were collected into new sterile microcentrifuge tubes for cell and tissue samples, respectively. Equal volumes of 70% ethanol were added to the collected samples, and the samples were vortex-mixed for 15 seconds. The RNA was further purified using the PureLink RNA Mini Kit according to the manufacturer’s instructions. The purity of the RNA was assessed using a Nanodrop, and the concentration was determined using Qubit™ RNA High Sensitivity (HS), Broad Range (BR), and Extended Range (XR) Assay Kits (Thermo, Cat #Q32852). The RNA was reverse-transcribed into cDNA using the Applied Biosystems High-Capacity RNA-to-cDNA Kit (Applied Biosystems, Cat #4387406). RT-qPCR was performed in a QuantStudio 7 Flex Real-time PCR System (Applied Biosystems). Most of the primers were ordered from Qiagen (Cat #330001) except for CLU exon-2 Forward: 5′-TCTCTGACAATGAGCTCCAAG-3′, exon-3 Reverse: 5′-TCCTCTAAACTGTTGAGCAAGG-3′; GAPDH: Forward: 5′-AGGTCGGTGTGAACGGATTTG-3′, Reverse: 5′-TGTAGACCATGTAGTTGAGGTCA-3′.

### Western immunoblotting

For the collection of cell medium fractions, the medium was obtained and centrifuged at 10,000 × g for 10 minutes at 4 °C. The resulting supernatant was collected and combined with protease and phosphatase inhibitors (PPI) (Thermo Fisher Scientific, Cat #78442). To prepare cell lysates, primary cells or IMG cells were washed with cold PBS and then lysed on ice. Neuronal protein extraction reagent (NPER, Cat #87792) with PPI was used for primary cells, while mammalian protein extraction reagent (MPER, Cat #7850) with PPI was used for IMG cells. The whole-cell lysate was centrifuged at 10,000 x g for 10 minutes at 4 °C, and the resulting supernatant was collected. For mouse cortical tissues, the Tissue Protein Extraction Reagent (T-PER) (Thermo Fisher Scientific, Cat #78510) containing PPI was used. Approximately 30 mg cortical tissue was homogenized using a Bullet Blender 24 homogenizer (Next Advance, Averill Park, NY) with 0.5 mm glass beads in 300 μl T-PER with PPI. The cortex lysate was obtained by centrifuging the sample at 10,000 x g for 8 minutes at 4 °C. Protein concentration was determined using a protein assay kit (Thermo Fisher Scientific, Cat #23225). The samples were then diluted in Laemmli sample buffer (Bio-Rad, Hercules, CA, USA) containing 2-mercaptoethanol (Bio-Rad) and boiled for 5 minutes at 95 °C. To separate the proteins, SDS-PAGE was performed, followed by transfer onto 0.2 μm pore-sized PVDF membranes (Bio-Rad). The membranes were blocked in 5% non-fat dry milk in TBS-T [100 mL of 10X TBS (200 mM Tris, 1.5 M NaCl), 900 mL of ddH_2_O, 10% of Tween-20, pH 7.6] for 1 hour at RT. Subsequently, the membranes were incubated overnight at 4 °C with primary antibodies diluted in TBS. The primary antibodies used were goat anti-CLU (1:2000, R&D, Cat #AF2747), goat anti-IL-1β (1:800, R&D, Cat #AF-401-NA), rabbit anti-CD11b (1:1000, Abcam, Cat #ab133357), and horseradish peroxidase (HRP) conjugated anti-β-actin (1:20,000, BioLegend, Cat #664804). After the overnight incubation, the membranes were washed three times with TBS-T and then incubated with the HRP-conjugated secondary antibody at RT for 1 hour. The secondary antibodies used were rabbit anti-goat HRP (1:3000, Invitrogen, Cat #31402) and goat anti-rabbit HRP (1:3000, Invitrogen, Cat #31462). The bands on the membrane were detected using a C-Digit Blot Scanner (LI-COR, Lincoln, NE, USA) after a 5-minute incubation with the enhanced chemiluminescence (ECL) reagent (BioRad, Cat #170-5061). Quantification was performed using Image Studio Version 4.0 imaging digitizing software and standardized to the internal loading control protein.

### Immunocytochemistry

Primary microglia and IMG cells were plated on 22 mm PDL-coated coverslips (Neuvitro, Cat #GG-22-PDL), individually placed in 35 mm cell culture dishes. After appropriate treatment, the cells were fixed with 2% paraformaldehyde (PFA) (Electron Microscopy Sciences, Cat #15710-S) for 15 minutes. The coverslips were then washed several times in PBS containing 0.1% saponin and blocked with serum corresponding to the host species of the secondary antibody. Next, the cells were incubated overnight at 4 °C with the primary antibodies diluted in blocking serum. The following day, the coverslips were thoroughly washed and incubated with the appropriate secondary antibodies for 2 hours at RT. Subsequently, the cells were incubated with 10 μM DAPI (Thermo, Cat #D1306) for 1 hour to stain the nuclei. After staining, the coverslips were mounted onto glass slides using mounting medium (Vector Labs, Cat #H1700). The slides were then imaged using a Zeiss Axio Observer inverted microscope. The primary antibodies used were rabbit anti-Iba1 (1:500, Wako, Cat #019-19741) and rat anti-CD11b (1:1000, Abcam, Cat #ab8878). The secondary antibodies used were donkey anti-rabbit (1:200, Invitrogen, Cat #A-31573) and donkey anti-rat (1:200, Invitrogen, Cat #A21208).

A skeleton analysis method on ImageJ (Fiji), as previously described (*76*), was used to quantify microglia morphology using immunofluorescent images of fixed primary microglia cells. The fluorescence images were converted to 8-bit. The Unsharp Mask feature under the Process tab was utilized to sharpen and enhance the image, while the noise de-speckling feature helped eliminate noise. Threshold adjustments were made to convert the image to binary. Subsequently, the AnalyzeSkeleton plugin was applied to skeletonize the image, and data on microglia process length were collected. The results were exported to an Excel spreadsheet. On the skeletonized image, several cell branches were measured using the line tool to determine the average length, and a consistent cutoff value was established across all analyzed images. The exported values were sorted by branch length from largest to smallest, and values above the cutoff were summed to calculate the total length of all branches collected from the image. This summed value was then divided by the number of cells in the corresponding image, yielding a normalized value used for further analysis.

### Luminex multiplex cytokine assay

Luminex cytokine assay was performed according to the manufacturer’s instructions (BioRad, Bioplex 200 Systems) using Bio-Plex Pro Reagent Kit (BioRad, Cat #12002798). Briefly, IMG cell lysates and media were collected following their respective treatments and mixed with PPIs to prevent protein degradation. 50 μL of the medium was added to 50 μL primary antibody-conjugated magnetic microspheres [BioRad, TNF-α (Cat #171G5023M) and IL6 (Cat #171G5007M)], and the mixture was incubated by shaking at 850 rpm for 30 minutes in a black 96-well plate at RT. After incubation with primary antibodies, each well was triple-washed with 1X wash buffer using a magnetic washer, then incubated with biotin-conjugated secondary antibodies by shaking at 850 rpm for 30 minutes at RT, triple-washed again, incubated with streptavidin/phycoerythrin by shaking at 850 rpm for 10 minutes, and washed two more times, finally suspended in 125 μL assay buffer by shaking at 850 rpm for 30 seconds at RT. Finally, the Bio-Plex reader was used to read the plate. The entire procedure was performed in a dark room.

### Generation of expression constructs for recombinant mouse CLU (rCLU)

The mouse *Clu* DNA was amplified by PCR from pCMV6-*Clu****_Exon2-9_*** plasmid (*25*) using the following primer pair: mouse *Clu*_NheI_Forward [5’-GCTG<u>GCTAGC</u>ATGAAGATTCTCCTG CTGTGCGTG-3’ (NheI restriction site is underlined)] and mouse *Clu*_AfeI_Reverse [5’-TCCA<u>AGCGCT</u>TTCCGC ACGGCTTTTCCTGCGGTA-3’ (AfeI restriction site is underlined)]. The mouse *Clu* DNA was then lifted out by PCR and subcloned into pcDNA3.1(-) expression vector harboring strep-tag II at the NheI and AfeI sites to generate pcDNA3.1(-)-*Clu****_Exon2-9_***-Strep II (mammalian expression vector for the mouse *Clu****_Exon2-9_*** with a C-terminal Strep II-tag) (Fig. S4). Sequences were verified by Sanger sequencing (Azenta, South Plainfield, NJ).

### Production and purification of rCLU

Recombinant mouse CLU was produced using FreeStyle 293-F cell system (Thermo Fisher Scientific). Briefly, FreeStyle 293-F cells were cultured in FreeStyle 293 expression medium at 37 °C with 8% CO_2_ on an orbital shaker at 125 rpm. Cells were transfected with pcDNA3.1(-)-mouseCLU-Strep II using 293fectin transfection reagent according to the manufacturer’s protocol. The culture medium was harvested at approximately 96 hours post-transfection, collected by centrifugation, and then concentrated using a 10 kDa molecular weight cutoff (MWCO) filter. The rCLU was then purified using a Strep-Tactin XT 4Flow high-capacity column (IBA, Cat #2-5011-005). SDS-PAGE and Western blot were used to examine protein purity. Pierce BCA protein assay kit was used to measure protein concentrations. Average yield was 30–35 mg of protein per liter of culture medium (Fig. S4).

### Y-maze two-trial test

The Y-maze two-trial test was conducted following previously described methods (*77*). A Y-maze apparatus with standard arm dimensions of 35 cm in length, 15.5 cm in height, and a lane width of 5 cm (Stoelting, Cat #60180) was utilized. Visual cues were consistently applied throughout the sessions. The testing environment was maintained at approximately 22 °C with subdued lighting (15-20 LUX) and minimal noise. Mice were acclimated to the behavior room for 1 hour prior to the test. The Y-maze two-trial test comprised two sessions: an acquisition trial and a retention trial, with a 30-minute interval. In the acquisition trial, one arm of the maze was blocked, allowing mice to explore the open arms for 5 minutes. After the interval, mice were reintroduced to the maze with all arms open for 5 minutes during the retention trial. To ensure consistency, mice were initially placed in the same corner of the maze, facing the wall. To prevent olfactory cues, the maze was cleaned thoroughly with 10% ethanol and Peroxigard between trials. The time spent in the novel arm was calculated as a percentage of the total time spent in the previously blocked arm during the first 30 seconds of the retention trial. An entry into an arm was recorded when 85% of the body was within the arm. Data tracking, recording, and analysis were performed using ANY-MAZE software (Stoelting, Wood Dale, IL). Mice with fewer than three arm entries (excluding the start arm entry) within the first minute of testing were identified as outliers and excluded from the analysis.

### Novel object recognition (NOR) test

The NOR test was conducted as previously described (*77*). The open field arena used in the test was a uniformly illuminated box measuring 40 x 40 x 35 cm (Stoelting, Cat #60101). NOR standard accessories from Stoelting Co. (Cat #62007s, Stoelting, Wood Dale, IL) were employed to ensure consistent features and prevent bias in mice preferences. The lighting conditions were adjusted to 15-20 LUX, RT was maintained at 22 °C, and efforts were made to minimize sound disturbance. Mice were allowed to acclimate in the behavior room for 1 hour prior to the test. The NOR test consisted of four sessions: habituation, familiarization, interval, and testing session. During the habituation phase, mice were allowed to explore and acclimate to the empty open field for 10 minutes. Twenty-four hours later, during the familiarization session, two identical objects were placed 5 cm from the wall in the open field. Each mouse was individually introduced into the open field, positioned facing away from the objects, and given 10 minutes to acclimate to the environment and the objects. Following a 45-minute interval, the mice were returned to the open field for the testing session, during which one of the familiar objects was replaced with a novel object. The mice were allowed 7 minutes to explore the objects, and the total time spent interacting with both objects throughout the testing session was recorded. The percentage of time spent with the novel object was calculated as the time spent interacting with it divided by the total interaction time with both objects during the entire test session. Between each phase of the test, the open field and objects were thoroughly cleaned with 10% ethanol and Peroxigard to eliminate olfactory cues. Data tracking, recording, and analysis were performed using ANY-MAZE software (Stoelting, Wood Dale, IL). Mice with a total object interaction time below 5 seconds were excluded from the analysis.

### Morris water maze (MWM) test

The MWM test was conducted following a previously described protocol with some adaptations (*78*). A circular pool measuring 109.2 cm in diameter (Hastings, Cat #60111S, RND 22, 4.2) was used, and spatial cues were consistently positioned around the pool and along the walls. The testing environment was set to 22°C, with low-intensity lighting (15-20 LUX) and minimal background noise. The water temperature was maintained at 24°C ± 2°C. The test comprised three stages: the visible stage, the hidden stage, and the probe trial. Each mouse completed Trial 1 before proceeding to subsequent trials. During Day 1 (visible stage - 5 trials/mouse), the platform was placed at different locations during each trial and positioned 1 cm above clear water. Mice were released from different quadrants and allowed to swim freely for 90 seconds or until they found the platform and stayed on it for at least 5 seconds. Mice that did not find the platform were gently guided to it and allowed to stay on it for 15 seconds. Days 2-4 involved a hidden training stage (5 trials/mouse/day), where the platform position remained constant (target quadrant) and was submerged 1 cm below opaque water. The water was made opaque by adding non-fat dry milk. Mice were released from different quadrants. On Day 5 (Test Day - 1 trial/mouse), the platform was removed, and one quadrant served as the starting location for all mice. Measurements recorded during the first 60 seconds of the testing trial included time spent and distance traveled in the target quadrant, total distance traveled, and average speed. The percentage of time and distance spent in the target quadrant was calculated by dividing the time or distance spent in the target quadrant by the total time or distance traveled across all quadrants. After each trial, the mice were transferred to a cage with paper towels under heating lamps for drying. The pool water was drained daily, and the pool was disinfected with 10% ethanol and Peroxigard to eliminate any olfactory cues. ANY-MAZE software (Stoelting, Wood Dale, IL) was used to track, record, and analyze data.

### MultiBrain sectioning

Mice were deeply anesthetized via intraperitoneal injection of ketamine (100 mg/kg body weight) and xylazine (10 mg/kg body weight) and transcardially perfused with PBS. The left-brain hemisphere was fixed overnight at 4 °C in 4% paraformaldehyde (Fisher, Cat #50-980-488), then washed and stored in PBS at 4 °C. The brains were then sent to NeuroScience Associates (NSA; Knoxville, TN), where they were sectioned using MultiBrain processing, as previously described (*79*). In brief, the hemispheres were treated with 20% glycerol and 2% DMSO to mitigate freeze artifacts, then embedded in a single gelatin block. After solidification, the embedded hemisphere block was quickly frozen by submersion in isopentane chilled to −70 °C using crushed dry ice and then mounted on the freezing stage of an AO 860 sliding microtome. The block of hemispheres was sectioned at a thickness of 35 µm in the coronal plane to encompass the hippocampus region (Bregma −1.0 to −4.0 mm). The sliced sections were sequentially collected into 12 containers containing an antigen preservation solution composed of 50% PBS, 50% ethylene glycol, and 1% polyvinyl pyrrolidone, with a pH of 7.0. After sectioning, each container held a serial set of one of every twelfth sections, forming composite sections that included individual sections from the hemispheres embedded in the block. This approach ensured uniform staining across the groups being compared.

### Campbell Switzer silver staining

The Campbell Switzer silver stain of plaque deposition was conducted by NeuroScience Associates as previously described (*80*). Briefly, the brain sections were removed from the storage cups and washed three times with distilled water (dH_2_0). They were then incubated with stirring in 2% ammonium hydroxide, followed by another wash in dH_2_O. Subsequently, the sections were placed in a silver-pyridine-carbonate solution and gently stirred for 40 minutes. Afterward, they were transferred to 1% citric acid for 3 minutes and then immersed in a pH 4.99 acetate buffer. The sections were developed by immersion in a freshly prepared physical developer composed of three different solutions over a light source: Solution A: dH_2_O, sodium carbonate; Solution B: dH_2_O, ammonium nitrate, silver nitrate, and tungstosilicic acid; Solution C: dH_2_O, ammonium nitrate, silver nitrate, tungstosilicic acid, and 37% formaldehyde. The development process was visually monitored and stopped by immersing the sections in a pH 4.99 acetate buffer, followed by a wash in dH_2_O and treatment with 0.5% sodium thiosulfate solution. After a thorough wash in dH_2_O, the sections were mounted and coverslipped.

### Aβ plaque imaging and quantification

Bright-field images of the stained sections were captured with a 10× objective on a Zeiss Axio Observer inverted microscope equipped with Zen Blue software. Montages were generated using the tile feature, which enabled selection of the start and end locations for image capture. Multiple images were acquired and stitched together to create a high-resolution composite image. Aβ plaque load analysis was performed on all stained sections using ImageJ software, following the protocol outlined on the NeuroScience Associates website (https://www.neuroscienceassociates.com/Alz-Plaque-Burden). Regions of interest (ROI) outlining the cortex-hippocampus regions were delineated using the elliptical tool, and separate traced images were created for analysis using the clear outside feature. The images were then converted to 8-bit, and the “Create Selection” and “Create Mask” options were utilized to add the ROI to the ROI manager. The total area of the ROI was measured from the traced image by clicking on “Measure”. Subsequently, the original traced images were reopened, converted to 8-bit, and thresholded (with lower and upper limits of 45 and 75, respectively) using Otsu’s method to accurately highlight Aβ plaques across sections. The area covered by plaques was measured by utilizing the “Create Selection” and “Create Mask” options to add the ROI to the ROI manager. Finally, the percentage of the area covered by plaques relative to the total cortex-hippocampus region was calculated for each image.

### Microglia Iba1 DAB staining

Free-floating mouse hemisphere sections were washed and incubated in PBS containing 3% hydrogen peroxide (H₂O₂) for 30 minutes at RT to quench endogenous peroxidase activity. After additional PBS washes, sections were blocked in a solution containing 3% normal donkey serum (NDS) and 0.3% Triton X-100 in PBS for 2 hours at RT to reduce non-specific binding. Sections were then incubated overnight at 4 °C with primary antibody, anti-Iba1 (1:1000, Wako), diluted in blocking solution. The following day, sections were washed in PBS containing 0.3% Triton X-100 and incubated for 1 hour at RT with a biotinylated secondary antibody, goat anti-rabbit IgG (1:500, Vector Laboratories). After washing, sections were incubated with the avidin-biotin complex (ABC) solution (Vector Laboratories), prepared 30 minutes prior to use (2 drops each of Reagent A and Reagent B per 5 mL blocking buffer), for 30 minutes at RT. Signal was developed using the DAB (3,3’-diaminobenzidine) substrate kit (Vector Laboratories) for approximately 3 minutes (2 drops Reagent 1, 4 drops Reagent 2, and 2 drops Reagent 3 per 5 mL of dH_2_O). Sections were then rinsed in PBS, mounted on glass slides, and cover-slipped with an antifade mounting medium (Vector Laboratories). Brightfield images were acquired using a Zeiss Axio Observer inverted microscope equipped with Zen Blue software. Imaging was performed at 10× and 20× magnifications.

### Single-nucleus RNA sequencing (snRNA-seq)

Mice were euthanized using CO_2_ inhalation, followed by the harvesting of cortical tissues, which were snap-frozen in dry ice to preserve RNA integrity. The frozen tissues were then transported to Novogene (Sacramento, CA) for snRNA-seq analysis. The process involved tissue dissociation to isolate single nuclei, followed by processing of the isolated nuclei for 10X Single Nucleus 3’ RNA Library Preparation. Subsequent steps included sequencing data analysis, beginning with transforming the original high-throughput sequencing image data into raw reads (referred to as Raw Data or Raw Reads). Sample demultiplexing was performed using 10-bp i5 and i7 sample index reads to generate paired-end Read1 and Read2 sequences, stored in FASTQ format. The snRNA-seq reads were processed using Cell Ranger software with default parameters, involving alignment to a reference, UMI collapsing, and initial quality control. Gene expression matrices containing cellular barcodes were filtered, and Seurat was used for data quality control. The Cell Ranger software employed the STAR aligner to perform splicing-aware alignment to the mouse genome, followed by transcript annotation and classification into exonic, intronic, and intergenic regions. Reads aligning to the transcriptome were mapped and subjected to UMI counting. Low-quality libraries were identified based on various criteria. Quality control metrics were calculated for each cell, including the total count and the proportion of mitochondrial gene counts. Cells failing any of these criteria were excluded from further analysis: less than 200 active genes per cell, genes with counts present in at most 3 cells, more than 50,000 total features (possibly indicating multiple cells combined), or over 50 % of counts coming from mitochondrial genes. Filtered feature matrices were loaded into Seurat for subsequent analysis, including normalization, feature selection to identify highly variable genes, and principal components analysis. Subsequently, shared nearest neighbor modularity optimization clustering, based on retained top principal components (typically 5 to 20), was used to identify transcriptionally related cell clusters. Dimensionality reduction techniques such as t-distributed stochastic neighbor embedding (t-SNE) and uniform manifold approximation and projection (UMAP) were employed to visualize cell clusters in 2D, providing insights into cell heterogeneity and relationships. Marker genes associated with specific cell types were identified by comparing differential expressions between clusters using statistical tests such as the Wilcoxon test. Enrichment analysis using the KEGG database identified significantly enriched pathways associated with DEGs. The degree of enrichment was measured using factors such as the Rich factor and q-value, with the top significantly enriched pathways displayed in the analysis report.

### Statistical Analysis

Statistical analyses were performed using GraphPad Prism version 6.01. Group comparisons were analyzed using one-way/two-way analysis of variance (ANOVA) followed by Tukey’s post hoc test or Student’s *t*-test, as appropriate. The data are presented as the group mean ± standard error of the mean (SEM). A *p*-value less than 0.05 was considered statistically significant.

## SUPPLEMENTAL MATERIALS

The PDF file includes:

Figs. S1 to S6

Table S1

## Acknowledgments

We thank the Transgenic and Gene Targeting Institutional Facility at the University of Kansas Medical Center for the generation of the novel *Clu*^−/−^ mouse model and the support of this Facility as a Shared Resource of the University of Kansas Cancer Center (NIH P30 CA168524). We would also like to thank the RIKEN BioResource Research Center for providing the *App*^NL-F/NL-F^ mouse model of AD under the Material Transfer Agreement.

## Author contributions

Conceptualization: L.Z. Data collection: P.R., H-J.M., V.N., S.K. Data analysis: P.R., H-J.M., V.N., S.K. L.Z. Writing—original draft: P.R., L.Z. Writing—review & editing: H-J.M., J.L.V., L.Z. Resources (Mouse Models): M.A.L., J.L.V., T.C.S. Funding acquisition: L.Z. Supervision: L.Z.

## Data and materials availability

All data needed to evaluate the conclusions in the paper are present in the paper and/or the Supplemental Materials.

