## Supplemental Figures Tables for "Clusterin regulates microglial inflammation and cognitive function independent of amyloid pathology in Alzheimer’s disease"

Punam Rawal *et al.*

**This PDF file includes:**

Figs. S1 to S6
Table S1

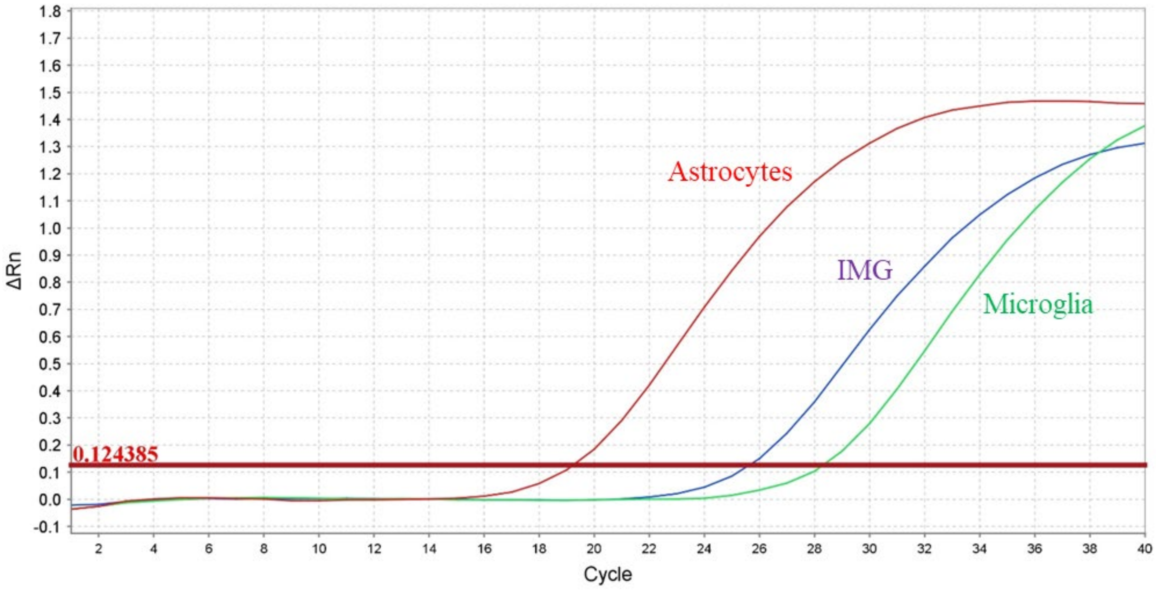


**Fig. S1.** Amplification plots from primary astrocytes, primary microglia, and IMG cells were analyzed for *Clu* mRNA expression by RT-qPCR.

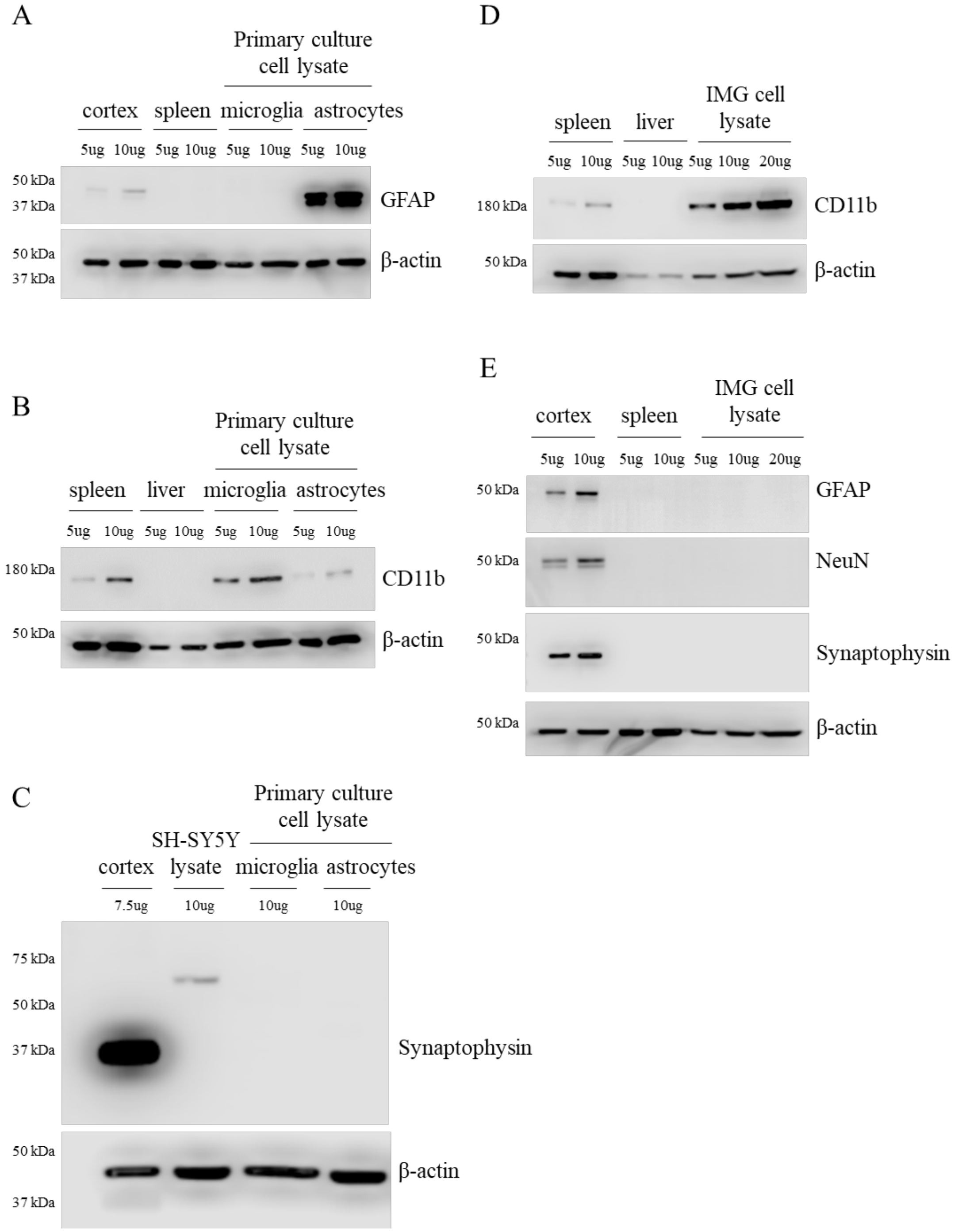


**Fig. S2.** Biochemical validation of cell models used in the study. **(A)** Primary mouse astrocytes are validated by a strong GFAP expression in the lysate. **(B)** Primary mouse microglia are validated by a strong CD11b expression in the lysate. **(C)** The culture specificity is confirmed by the absence of the neuronal marker synaptophysin in both primary astrocyte and microglia lysates. **(D)** IMG cells are validated by a strong CD11b expression in the lysate. **(E)** IMG cell culture specificity is confirmed by the absence of the astrocytic marker GFAP and neuronal markers NeuN and synaptophysin in cell lysates.

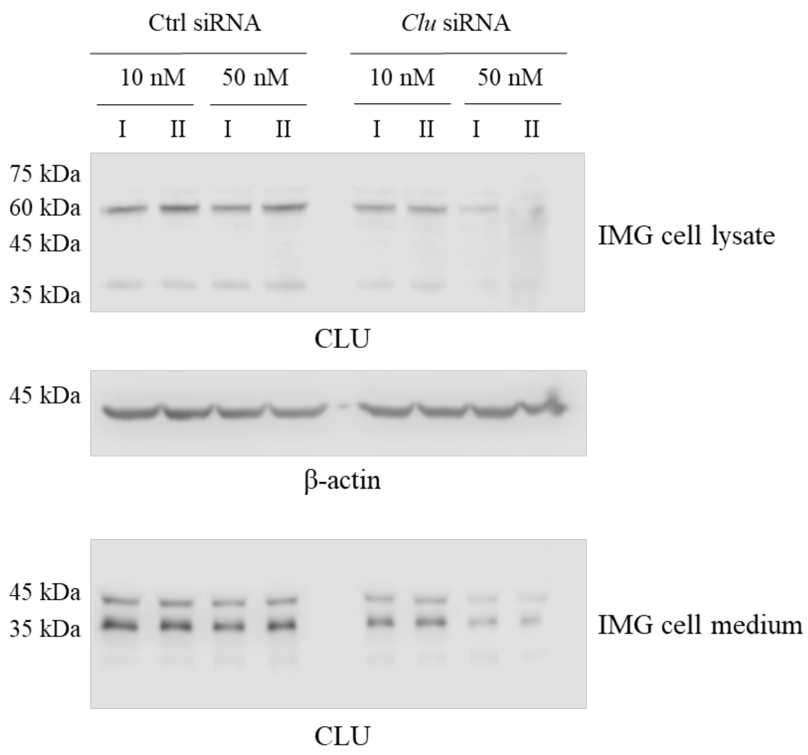


**Fig. S3.** Assessment of *Clu* siRNA transfection efficiency. IMG cells were transiently transfected with 10 nM or 50 nM *Clu* siRNA or a scramble control. Transfection with 50 nM *Clu* siRNA significantly reduced CLU expression in both cell lysate and culture medium.

A

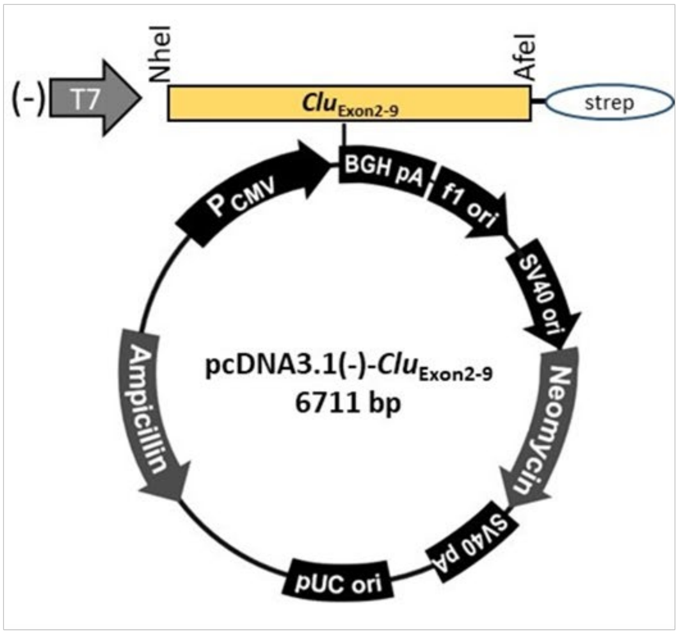


B

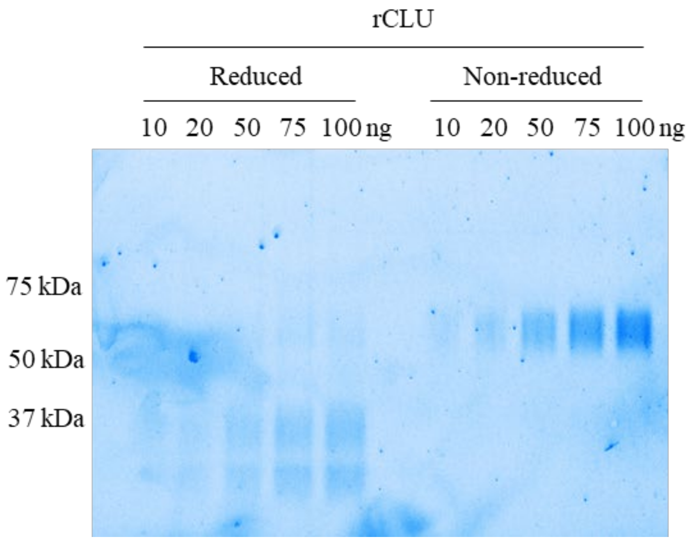


**Fig. S4.** Generation and validation of recombinant mouse mature CLU (rCLU). **(A)** A mammalian expression vector for mouse *Clu* (containing Exon2-9) with a C-terminal Strep II-tag was generated and used to produce recombinant mouse mature CLU. **(B)** The resulting rCLU was purified using Strep-Tactin XT 4Flow high-capacity columns, and purity was confirmed by TGX Stain-Free FastCast polyacrylamide gels under both reducing and non-reducing conditions. Under reducing conditions, rCLUα and rCLUβ subunits were detected at ~34 kDa and ~37 kDa. Under non-reducing conditions, the full-length rCLU appeared as a single band at ~70 kDa.

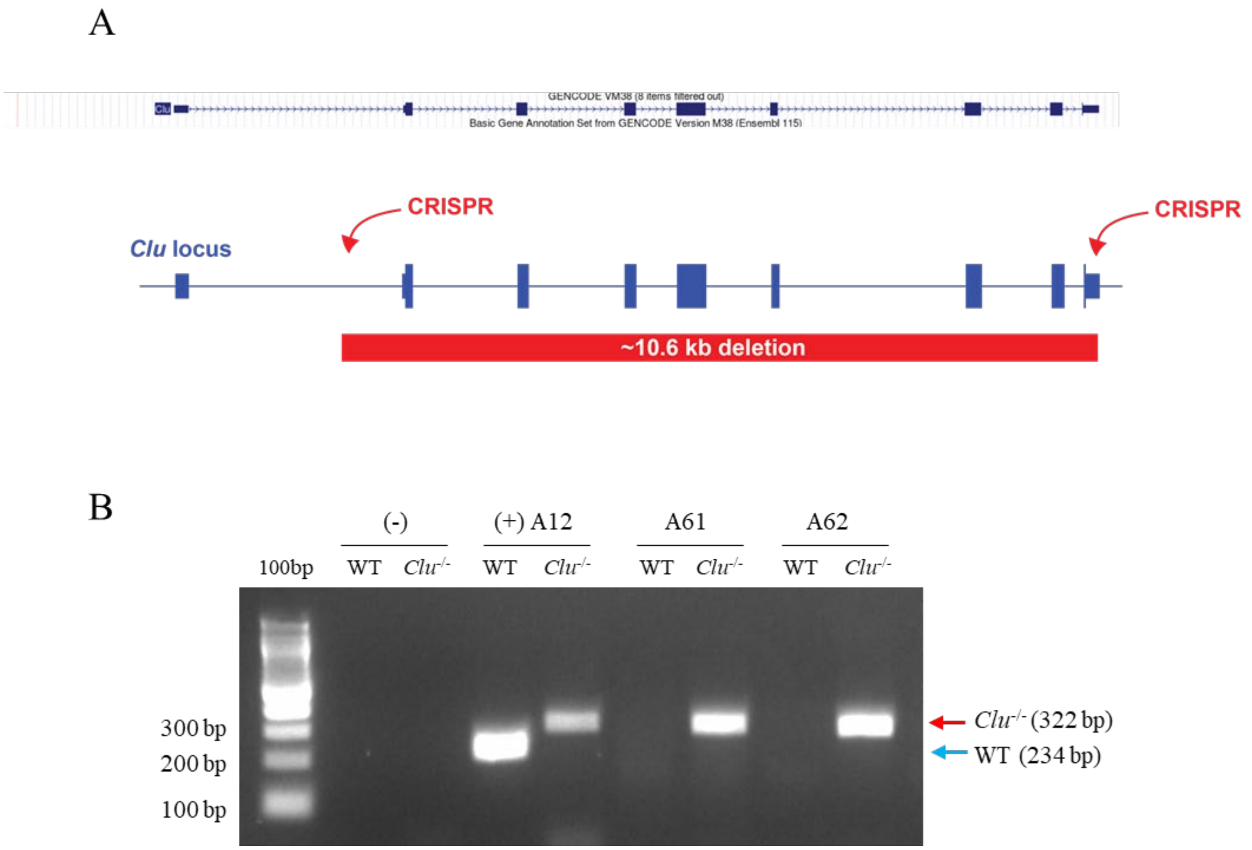


**Fig. S5.** Generation and validation of novel *Clu*$^{-/-}$ mouse model. **(A)** Schematic depiction of the total knockout mouse model of *Clu*. The *Clu* gene, situated on chromosome 14 in the mouse genome, underwent CRISPR deletion spanning 10,628 base pairs (from nucleotide position 3,024 to 13,651). This deletion starts between exons 1 and 2 and extends to the near end of exon 9, resulting in the complete removal of the *Clu* gene in the mouse genome. **(B)** Representative genotyping results for WT and *Clu*$^{-/-}$ mouse models showing PCR products from WT at 234 bp and CLU$^{-/-}$ at 322 bp.

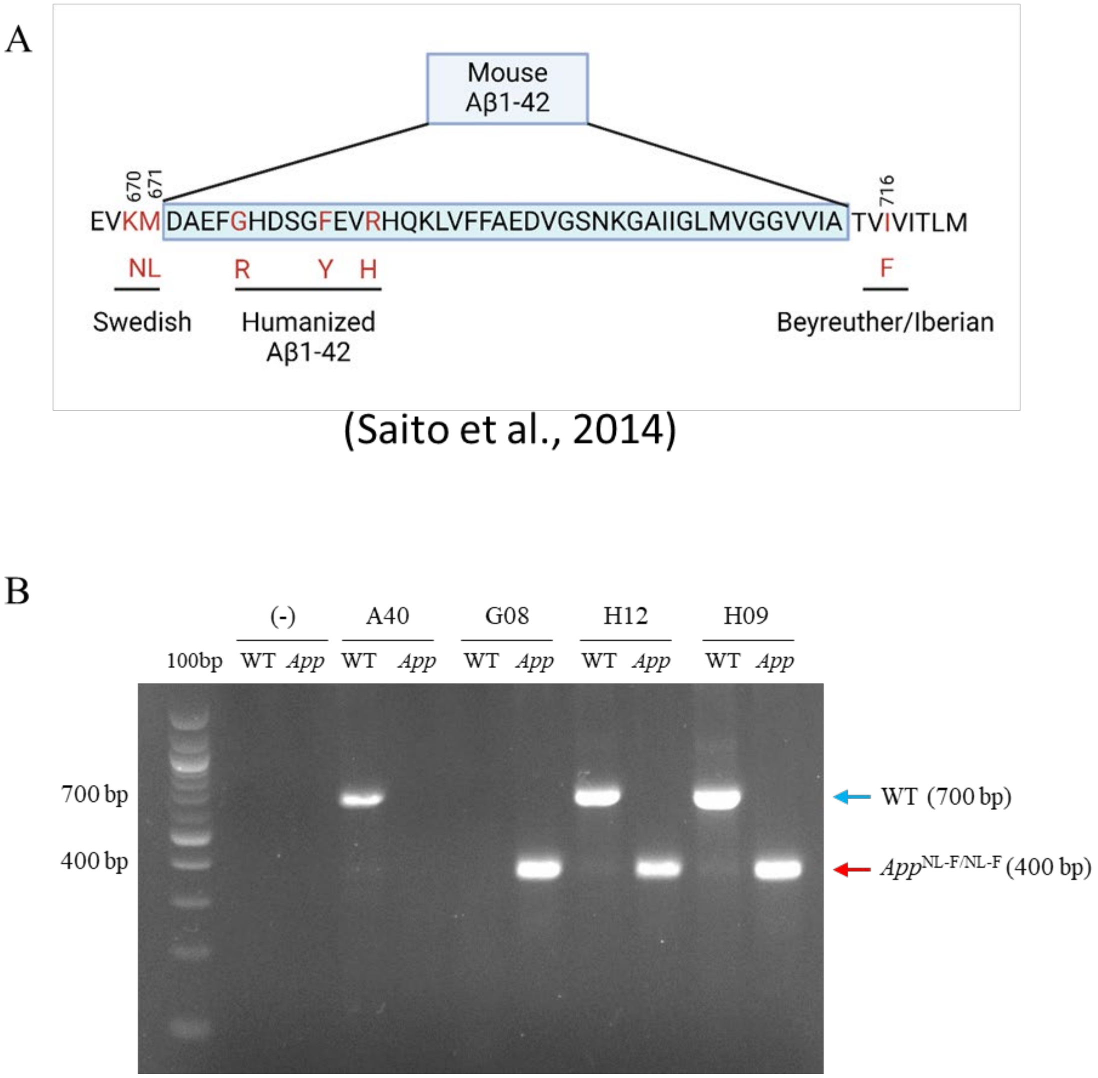


**Fig. S6.** Validation of *App*$^{NL\text{-}F/NL\text{-}F}$ mouse model. **(A)** Schematic representation of the alterations to the mouse APP in the *App*$^{NL\text{-}F/NL\text{-}F}$ mouse model. The mouse Aβ sequence was humanized by substituting three amino acids, followed by the introduction of the Swedish and Beyreuther/Iberian mutations via knock-in technology. Adapted from Saito *et al.* (*Nat Neurosci* 17(5):661-3, 2014) and created with BioRender. **(B)** Representative genotyping results for WT and *App*$^{NL\text{-}F/NL\text{-}F}$ mouse models showing PCR products from WT at 700 bp and *App*$^{NL\text{-}F/NL\text{-}F}$ at 400 bp**.**

**Table S1. Primer sequences used in the genotyping of mouse models.**

| Gene | Primer | Sequence | Size |
|---|---|---|---|
| *Clu*$^{-/-}$ | WT_F | 5'-CTTTTCCCTGTTCCTGTGGGT-3' | 234 bp for WT |
| | WT_R | 5'-TGAGGCAGGGCAGATGAATTT-3' | |
| | *Clu*$^{-/-}$_F | 5'-AGTCTCTTCTCCCACACCACT-3' | 322 bp for *Clu*$^{-/-}$ |
| | *Clu*$^{-/-}$_R | 5'-CAGTGTTGTAACGGGAAGGGAA-3' | |
| *App*$^{NL-F/NL-F}$ | E16WT | 5'-ATCTCGGAAGTGAAGATG-3' | 700 bp for WT |
| | WT | 5'-TGTAGATGAGAACTTAAC-3' | |
| | E16MT | 5'-ATCTCGGAAGTGAATCTA-3' | 400 bp for *App*$^{NL-F/NL-F}$ |
| | LoxP | 5'-CGTATAATGTATGCTATACGAAG-3' | |
